# L-norvaline reverses cognitive decline and synaptic loss in a murine model of Alzheimer’s disease

**DOI:** 10.1101/354290

**Authors:** Baruh Polis, Kolluru D Srikanth, Evan Elliott, Hava Gil-Henn, Abraham O. Samson

## Abstract

The urea cycle plays a role in the pathogenesis of Alzheimer’s disease (AD). Arginase-I accumulation at sites of amyloid-beta deposition is associated with L-arginine deprivation and neurodegeneration. Moreover, a positive interaction between the arginase-II and mTOR-S6K1 pathways promote inflammation and oxidative stress. In this study, we treated 3xTg-AD mice exhibiting increased ribosomal protein S6 kinase beta-1 (S6K1) activity and wild-type (WT) mice with L-norvaline. The substance combines unique properties of arginase and S6K1 inhibition. The treated 3xTg-AD mice demonstrated significantly improved acquisition of spatial memory, associated with a substantial reduction of microgliosis and shift from activated to resting phenotype. Moreover, an increase of dendritic spines density and escalation of the expression rates of neuroplasticity related proteins were followed by a decline in the levels of amyloid-beta toxic oligomeric and fibrillar species in the hippocampus. Our findings associate local amyloid-beta-driven and immune-mediated <u>response</u> with altered L-arginine metabolism and suggest that arginase and S6K1 inhibition by L-norvaline can delay progression of AD.

## Introduction

Alzheimer’s disease (AD) is a slowly progressive neurodegenerative disorder with an insidious onset. Advanced age is a prominent risk factor of AD and other metabolic diseases, such as type II diabetes and atherosclerosis. Their causal mechanisms are multifaceted and not fully interpreted (Kandimalla, Thirumala, and Reddy 2016). Recent clinical and experimental data demonstrate that neurodegenerative disorders and other metabolic diseases often display a coexisting metabolic dysfunction, which may exacerbate neurological symptoms (H. Cai et al. 2012). Moreover, it has been suggested that nutritional behavior is a key triggering factor in the pre-symptomatic stages of AD (Ríos et al. 2014). Additional hypothesis emphasizing impairment of bioenergetic metabolism as a key contributing mechanism to the pathogenesis of AD has been proposed recently (Sonntag et al. 2017). According to the authors, AD is characterized by a combination of several interrelating pathological events that include bio-energetic, metabolic, neurovascular, and inflammatory processes. Other researchers point out the fact that brain hypo-metabolism occurs decades prior to the clinical manifestation, which suggests that metabolic dysfunction is an important contributing factor to AD (Wilkins and Trushina 2018). Until now, the lack of comprehensive analysis and particular methodology has limited the development of AD diagnosis and therapy.

New unbiased lipidomics and metabolomics approaches were applied to analyze the changes in post-mortem brains from patients suffering from AD (Paglia et al. 2016). Several significantly altered metabolites that distinguished AD from controls were detected. L-arginine metabolic pathway was identified and mentioned as highly implicated in the development of AD. Similar results indicating the dysregulation of L-arginine metabolism have been acquired in a rodent model of AD (J. Yu et al. 2017). Hence, the brain’s urea cycle is under intensive investigation.

The brain is identified as an important metabolic reactor, which is actively involved in the urea cycle (Gropman 2010). Several diseases were indicated as having high brain urea concentrations and misbalanced expression of the related genes. For example, it was reported that brains of patients suffering from Huntington’s disease (HD) contain about threefold more urea compared to the healthy controls (Patassini et al. 2015). These findings have been proved in a transgenic sheep model of HD (Handley et al. 2017) Additionally, the data indicating that the postmortem brains of AD patients contain significantly higher concentrations of urea compared to the control group have been published recently (Xu et al. 2016). Consequently, the hypothesis linking build-up of urea in the brain to toxic levels causing brain damage and finally dementia was proposed (Handley et al. 2017).

Recent studies suggest a role of arginase as a key element of the urea cycle, converting L-arginine to urea and L-ornithine, in AD, cardiovascular and other metabolic diseases (Pernow and Jung 2013). Significantly decreased levels of L-arginine and L-ornithine in the cortices of AD patients have been observed (Gueli and Taibi 2013). Additionally, the activity and expression level of nitric oxide synthase (NOS) and arginase are meaningfully altered in AD brains in a region-specific manner. Namely, the activity of arginase is significantly higher and NOS is lower in the hippocampi of AD patients (Liu et al. 2014). Moreover, there are indications that arginase-II (ARGII) is mostly responsible for overall increased arginase activity in the brain.

Arginase is ubiquitous in early life forms and currently existing phyla (Dzik 2014). Plants, bacteria, and yeasts have a single form of arginase (ARGII) localized in mitochondria. Vertebrates process nitrogen by another cytosolic isoform of arginase (ARGI) (Caldwell et al. 2015). Different genes encode these two isoenzymes, which demonstrate distinctive cell and tissue patterns of distribution. In humans, ARGI is generally presented as a cytosolic enzyme of the liver, and also expressed in various regions of the brain. ARGII, described in the literature as a kidney-type, and is ubiquitously expressed at a low level within the mitochondria of various organs (Lange et al. 2004).

The presence of both ARGI and ARGII in the brain, especially in hippocampal neurons, was confirmed by immunocytochemical analysis of rat brain sections (Peters et al. 2013). Moreover, it was established that ARGII is the predominant isoform of the human frontal cortex (Morris, Bhamidipati, and Kepka-Lenhart 1997). The expression of two isoforms of arginase can be impelled in different tissues via exposure to a variety of cytokines and catecholamines (Sidney and Morris 1992), (Lange et al. 2004). ARGII is inducible by many stimuli including lipopolysaccharide (LPS), tumor necrosis factor alpha (TNFα), oxidized low-density lipoprotein (LDL), and hypoxia (Ryoo et al. 2011). Moreover, it was demonstrated that activation of ARGII is associated with its translocation from the mitochondria to the cytosol (Pandey et al. 2014).

Arginase was shown to be a neuro-protective factor and to support neuro-regeneration *in vitro* (Esch et al. 1998), (Deng et al. 2009). Nevertheless, upregulation of arginase has been associated with several diseases of the neural system, such as AD, Parkinson’s disease (PD), multiple sclerosis, stroke, traumatic brain injury and several retinal diseases (Caldwell et al. 2015), (Shosha et al. 2016). In addition, there are data demonstrating augmented ARGII gene expression in AD brains, and a potential association of the gene with the risk of developing AD has been hypothesized (Hansmannel et al. 2010). Furthermore, it was demonstrated that ARGII deficiency reduces the rate of hyperoxia-mediated retinal neurodegeneration (Narayanan et al. 2014), suggesting involvement of arginase in the pathogenesis of neuronal degeneration via excessive activation of the excitotoxic NMDA receptors (Pernet, Bourgeois, and Polo 2007). At the molecular level, a mutual positive regulation between ribosomal protein S6 kinase beta-1 (S6K1) and ARGII in endothelial inflammation processes and aging was demonstrated (Yepuri et al. 2012). So, it was suggested that ARGII and mTOR-S6K1 pathways play an important role in promoting oxidative stress, inflammation, senescence and death, which eventually leads to metabolic disorders (X.-F. 2013). Therefore, targeting ARGII was recently proposed as a means of decelerating age-related diseases treatment (Xiong et al. 2017).

In addition to deviation of arginase levels, overexpression of the gene responsible for ornithine transcarbamylase, another enzyme of the urea cycle, was identified in the AD brain (Bensemain et al. 2009), indicating a complex anomaly of L-arginine metabolism in AD. Subsequently, L-arginine and its derivatives were trialed with patients suffering from various neurological disorders. L-arginine administration within 30 minutes of stroke was shown to significantly decrease frequency and severity of stroke-like symptoms (Koga et al. 2005). Supplement of 1.6 g/day of L-arginine for three months in patients with senile dementia was found to increase cognitive function by about 40% (Ohtsuka and Nakaya 2000), and our unpublished results suggest the same is true for patients suffering from AD.

In healthy individuals, L-arginine is transported from the circulating blood into the brain via Na+-independent cationic amino acid transporter (CAT1) expressed at the blood–brain barrier (BBB) (O’Kane et al. 2006). The influx transport of the amino acid at the rat BBB is saturable with a Michaelis-Menten constant (Km) value of 56 μM. The physiological serum concentration of L-arginine is significantly higher in rodents (about 170 μM) and humans (about 100 μM) (Stoll, Wadhwani, and Smith 1993). Since L-arginine in mammals is derived mostly from renal *de novo* synthesis and dietary intake, CAT1 at the BBB functions as a supply pathway for L-arginine to the brain (Tachikawa and Hosoya 2011). So, despite the capability of this amino acid to pass the BBB, the capacity of its transporter is very limited (Shin et al. 1985), which makes oral administration of arginine insufficient to show all of its possible neurotrophic properties. Consequently, a hypothesis linking L-arginine brain deprivation and the development of AD cognitive deficiency was proposed (Kan et al. 2015). The theory has found an experimental approval. Fonar *et al.* bypassed the BBB by chronic intraventricular administration of the amino acid in mice (Fonar et al. 2018). The treatment improved significantly spatial memory acquisition in 3xTg-AD mice via a reduction of oxidative stress and apoptosis.

L-arginine is a mutual substrate both for arginase and for nitric oxide synthase (NOS) as well, which produce urea, and nitric oxide (NO) with L-citrulline respectively (Nelson and Cox 2004). So, the bioavailability of L-arginine is a regulating factor for NO synthesis; thus, its production is seriously dependent upon arginase activity, which easily depletes the substrate (Balez and Ooi 2016). There are data showing a serious reduction of NOS activity in AD brains, with decrease in the levels of NOS1 (neuronal) and NOS3 (endothelial) proteins (Liu et al. 2014). Venturini *et al.* demonstrated that amyloid peptides strongly inhibit the NOS1 and NOS3 activity in cell-free and cellular systems, which provides a novel molecular mechanism for AD development (Venturini et al. 2002).

A growing body of evidence indicates that NO possesses neuroprotective properties. NO triggers vasodilation and increases blood supply to neurons, which reduces their susceptibility to oxidative stress (Durante, Johnson, and Johnson 2007). Moreover, NO regulates Ca^2+^ influx into the neurons via inhibition of NMDA receptors at glutamatergic synapses, protecting the cells from overstimulation and, subsequently, excitotoxicity (Ditlevsen et al. 2007). Additionally, L-Citrulline, which is a byproduct of the reaction, catalyzed by NOS, was shown to possess neuroprotective properties (Yabuki et al. 2013).

To investigate the role of NO in the development of AD the APP/NOS2^−/−^ mice were designed (Wilcock et al. 2008). The mice express the human Swedish mutation (APPSw; Tg2576) on a homozygous mouse NOS2 knockout background, and exhibit behavioral deficits and biochemical hallmarks of AD during aging. Working with this model of AD Kan *et al.* demonstrated that ARGI is highly expressed in regions of Aβ accumulation (Kan et al. 2015). It was proved that pharmacologic disruption of the arginine utilization pathway by irreversible inhibition of ornithine decarboxylase (ODC) with α-difluoromethylornithine (DFMO) protects the mice from AD-like pathology and reverses memory loss. The authors suggest that L-arginine depletion is responsible for neuronal cell death and cognitive deficits in the course of AD development.

Another knockout (NOS3 ^−/−^) mice model was shown to develop Alzheimer’s like pathology (Austin et al. 2013). Moreover, a negative correlation between capillary expressed NOS3 and Alzheimer’s lesion burden was reported (Jeynes and Provias 2009). We hypothesize that upregulation of arginase activity and consequent L-arginine and NO deficiency in the brain lead to the manifestation of AD pathology. Therefore, we propose to target arginase to ameliorate the symptoms of the disease. We use an arginase inhibitor L-norvaline, which acts via negative feedback regulation. It was proved that arginase inhibition by L-norvaline amplifies the rate of NO production and reduces urea production (Chang, Liao, and Kuo 1998). L-norvaline has been used successfully to treat artificial metabolic syndrome in a rat model (El-Bassossy et al. 2013). Moreover, there are reports showing that L-norvaline inhibits S6K1 as well and possesses anti-inflammatory properties (Ming et al. 2009), so the application of the drug could be extremely effective in AD.

In this study, we used 3xTg-AD mice and treated them with L-norvaline dissolved in water. Animals from the control group displayed significant memory deficiency compared to the experimental group, evident in two paradigms. The observed cognitive effect was associated with reduced rates of **β**-amyloidosis, microgliosis, and astrodegeneration. Moreover, the treatment reversed dendritic spine deficiency in the cortices and hippocampi and amplified the expression rate of pre- and postsynaptic proteins.

## Results

### L-norvaline ameliorates memory deficits in 3xTg-AD mice

To investigate whether L-norvaline treatment affects learning and memory in AD pathogenesis, a set of behavioral tests was performed after the mice received two-month-long treatment starting at the age of 4 months.

Short-term working memory was assessed in the Y-maze spontaneous alternation test. A significant effect of the treatment upon “percentage of alternations” was revealed by a One-Way ANOVA test (F(3,54)=4.525, p=0.0067). Post-hoc Tukey’s Multiple Comparison Test indicated that control 3xTg-AD mice showed a lower percentage of alternation than L-norvaline treated mice (p<0.05) (Fig. 1a). L-norvaline treatment had no significant effect on alternative behavior of the non-transgenic (non-Tg) mice. The total distance moved by the animals during the test was not affected by the treatment (p=0.86) (Fig. 1b).

Morris Water Maze (MWM) was used to determine the effect of L-norvaline upon spatial memory. In the visible platform test there were no significant differences between the groups, (p=0.99), which indicate that L-norvaline treatment had no influence upon the mice motility or vision (Fig. 1c).

In the hidden platform-swimming test there were significant differences between the groups revealed with repeated measures ANOVA F(3,12) = 7.725, p=0.0039. Tukey’s Multiple Comparison test detected significant (p<0.01) effect of the treatment upon memory acquisition in 3xTg-AD group (Fig. 1d).

In the probe trial on the last day of testing, the platform was removed. The relative time spent by mice in the target quadrant, where the hidden platform was previously placed, was significantly greater within the L-norvaline treated group compared to the control group (34.8 ± 8.06 and 24.5 ± 7.88; p<0.05; Fig. 1e). The swim speed of the animals was not affected by the treatment (Fig. 1f). The overall results of the behavioral experiments suggest that L-norvaline improves spatial memory acquisition in 3xTg-AD mice.

**Figure 1.**
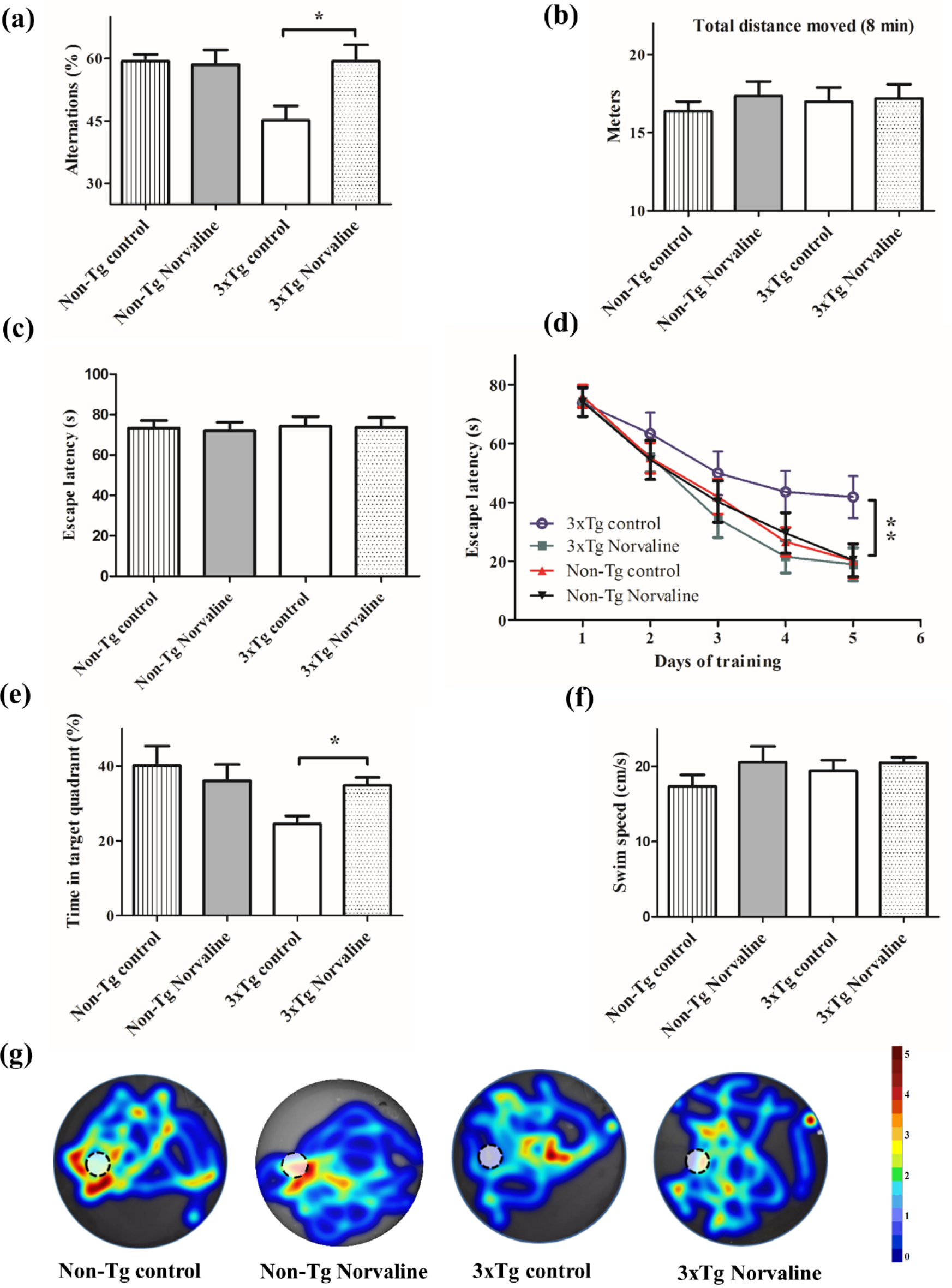
L-norvaline effects on behavior. **Y-maze** spontaneous alternation test. (a) Percentage of alternation. (b) Total distance moved during the trial. **MWM** (c) Visible platform test (averaged escape latencies of two trials). (d) Hidden platform test (averaged escape latencies of two trials/day). (e) Time spent by mice in target quadrant in probe trial. (f) Average swimming speed on the day with probe test. (g) Heat maps summary of search intensity during probe trials, where the platform is indicated by a dashed circle. High dwell time across the pool area is indicated by colors close to red, whereas colors close to blue indicate lower dwell time (arbitrary scale). Data are shown as means ± SEM. (n = 15 for non-TG groups, n=14 for 3xTg-AD groups). *p<0.05,** p < 0.01.

### L-norvaline treatment reduces the quantities of Aβ fibrils and prefibrillar oligomers in the hippocampi of 3xTg-AD mice

Recent evidence suggests the role of soluble amyloid oligomeric and fibrillar species in synaptic dysfunction, neuronal apoptosis and brain damage (Gadad, Britton, and Rao 2011). To investigate the quantities of toxic prefibrillar Aβ oligomers and fibrillary forms we examined the A11 and OC immunoreactivity of the hippocampal lysates. Pooled equal amounts (10 μg) of total lysates from the hippocampi of each group of mice were run on SDS polyacrylamide gels and incubated with appropriate antibodies (Supplementary Table 1 and Supplementary Fig. 2). Treatment with L-norvaline resulted in a substantial (about 30%) reduction in the levels of A11-reactive oligomers and OC-reactive fibrillar species (Table 1).

**Table 1.** L-norvaline reduces the level of fibrillar amyloid and pre-fibrillar oligomers in the hippocampal brain lysates of 3xTg-AD mice (blots summary). Western blot summary with β-actin normalized trace quantities of Aβ fibrils (a) and prefibrillar oligomers (b).

| Actin Normalized Trace Quantity |  |  |  |
| --- | --- | --- | --- |
| MW (kDa) | Control | Norvaline | Change (%) |
| 135.938 | 514 | 265 |  |
| 89.692 | 428 | 518 |  |
| 79.654 | 574 | 308 |  |
| 68.494 | 590 | 521 |  |
| 62.231 | 725 | 593 |  |
| 50.635 | 1703 | 1206 | ~30% |
| 43.94 | 1799 | 1185 |  |
| 37.763 | 1150 | 1289 |  |
| 25.766 | 4254 | 3229 |  |
| **Sum** | **11737** | **9114** | **~22%** |

**Table 1. L-norvaline reduces the level of fibrillar amyloid and pre-fibrillar oligomers in the hippocampal brain lysates of 3xTg-AD mice (blots summary).** Western blot summary with β-actin normalized trace quantities of Aβ fibrils (a) and prefibrillar oligomers (b).
| Actin Normalized Trace Quantity |  |  |  |  |
| --- | --- | --- | --- | --- |
| MW (kDa) | Control | Norvaline | Change (%) | Structure |
| 85.776 | 787 | 141 |  |  |
| 61.237 | 1622 | 1288 |  |  |
| 48.587 | 8723 | 6127 | ~30% | **Dodecamer** |
| 40.324 | 150 | 30 | ~80% | **Octamers** |
| 36.076 | 393 | 456 |  |  |
| 33.865 | 406 | 462 |  |  |
| 27.314 | 2784 | 1502 | ~46% | **Hexamers** |
| 24.216 | 1218 | 1245 |  |  |
| **Sum** | **16084** | **11251** | **~30%** |  |

### L-norvaline decreases the rate of beta-amyloidosis in the cortex of 3xTg-AD mice

To examine the effect of the treatment upon total amyloid burden in the brains of 3xTg-AD mice coronal brain sections were stained with human APP/Aβ-specific antibody. By six months of age, 3xTg-AD mice exhibited enhanced intracellular deposition of Aβ in layer IV-V of the PFC (Fig. 2a-c), hippocampus (Fig. 2d-f,i), and amygdala (Fig. 2g-h).

**Figure 2.**
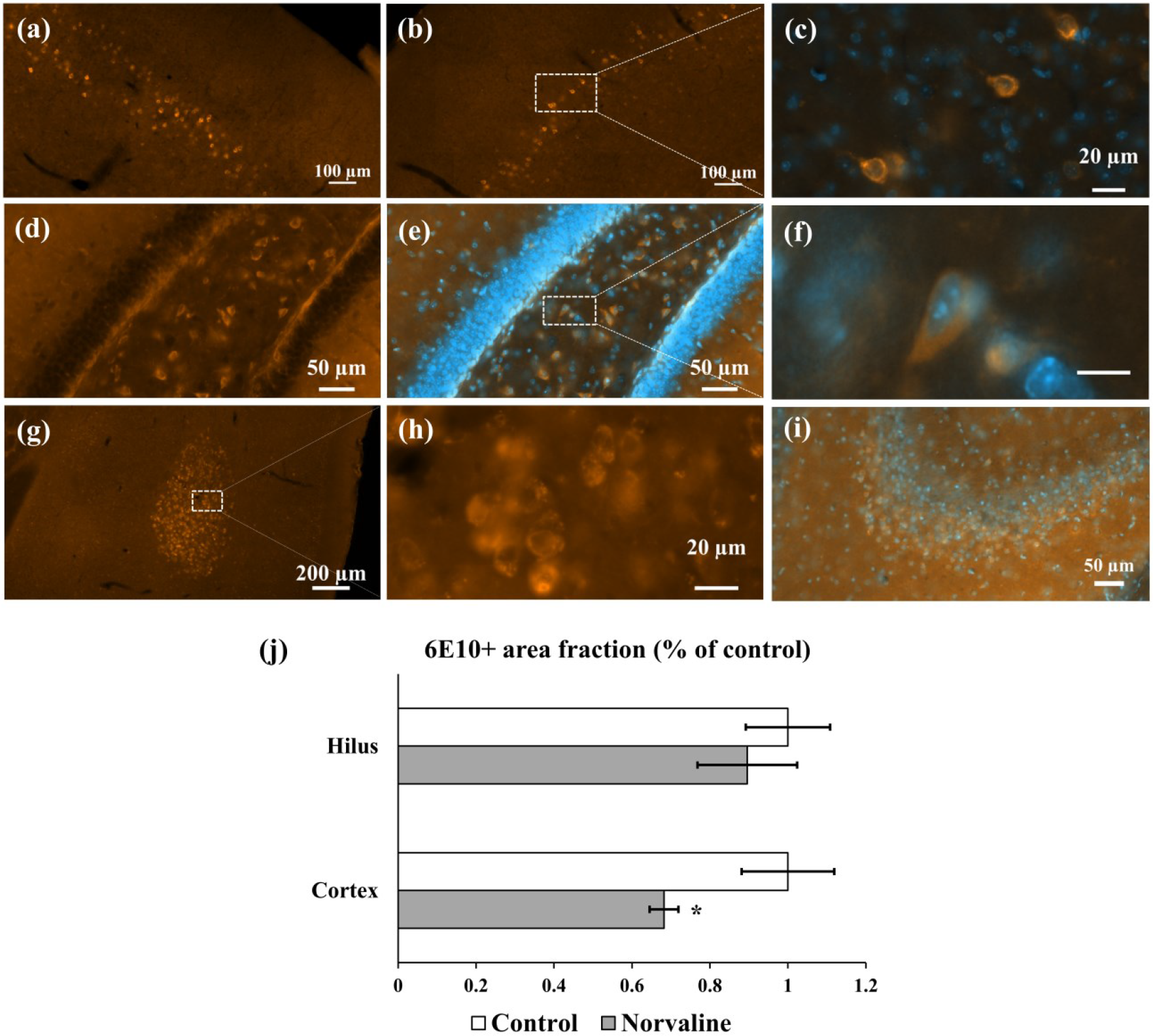
L-norvaline effect on total amyloid burden. Quantification of the area occupied by Aβ immunoreactivity in brains of 3xTg-AD mice receiving vehicle (control) or L-norvaline. A significant effect of the treatment on the rate of 6E10 immunoreactivity in PFC (a) control, (b) treatment (p=0.032). The insets represent 100× magnification of double staining with DAPI. Intense Aβ deposition in the dentate gyrus (d-f)) (f scale bar-20 μm), amygdala (g-h), and CA3 area (i). Plot of the immunoreactive area fraction for hilus and PFC (j). Student’s t-test was used to compare means between two groups, *p<0.05 (n=15, three mice per group).

We did not detect significant differences in the rate of Aβ deposition between the two groups of the animals in hippocampi; however, general trend was in slight reduction of the 6E10 positivity in the treated group. Nevertheless, we observed a significant (p<0.05) reduction in the rate of 6E10 positivity in the cortex of 3xTg mice treated with L-norvaline. The data indicate effect of L-norvaline on the rate of β-amyloidosis in 3xTg-AD mice.

### L-norvaline increases pyramidal cells dendritic spine density in the cortex and the hippocampus of 3xTg-AD mice

Dendritic spines are specialized structures on neuronal processes, which are involved in neuronal plasticity (Frankfurt and Luine 2015). A strong correlation between dendritic spine density and memory acquisition was demonstrated in rodents using different behavioral paradigms (Jedlicka et al. 2008). In the present study, dendritic spines were examined by Golgi staining and a comparison was made between vehicle- and L-norvaline-treated groups. Spine density was quantified on secondary basal dendritic segments located farther than 40 μm from the soma of the layer III pyramidal cells and secondary oblique dendritic branch localized in the stratum radiatum of the CA1 hippocampal pyramidal neurons. The analysis revealed a significant effect of the treatment on dendritic spine density. In the cortex of the treated mice the average increase was 18.6% (p=0.019), and 19.8% (p=0.012) in the hippocampus.

**Figure 3.**
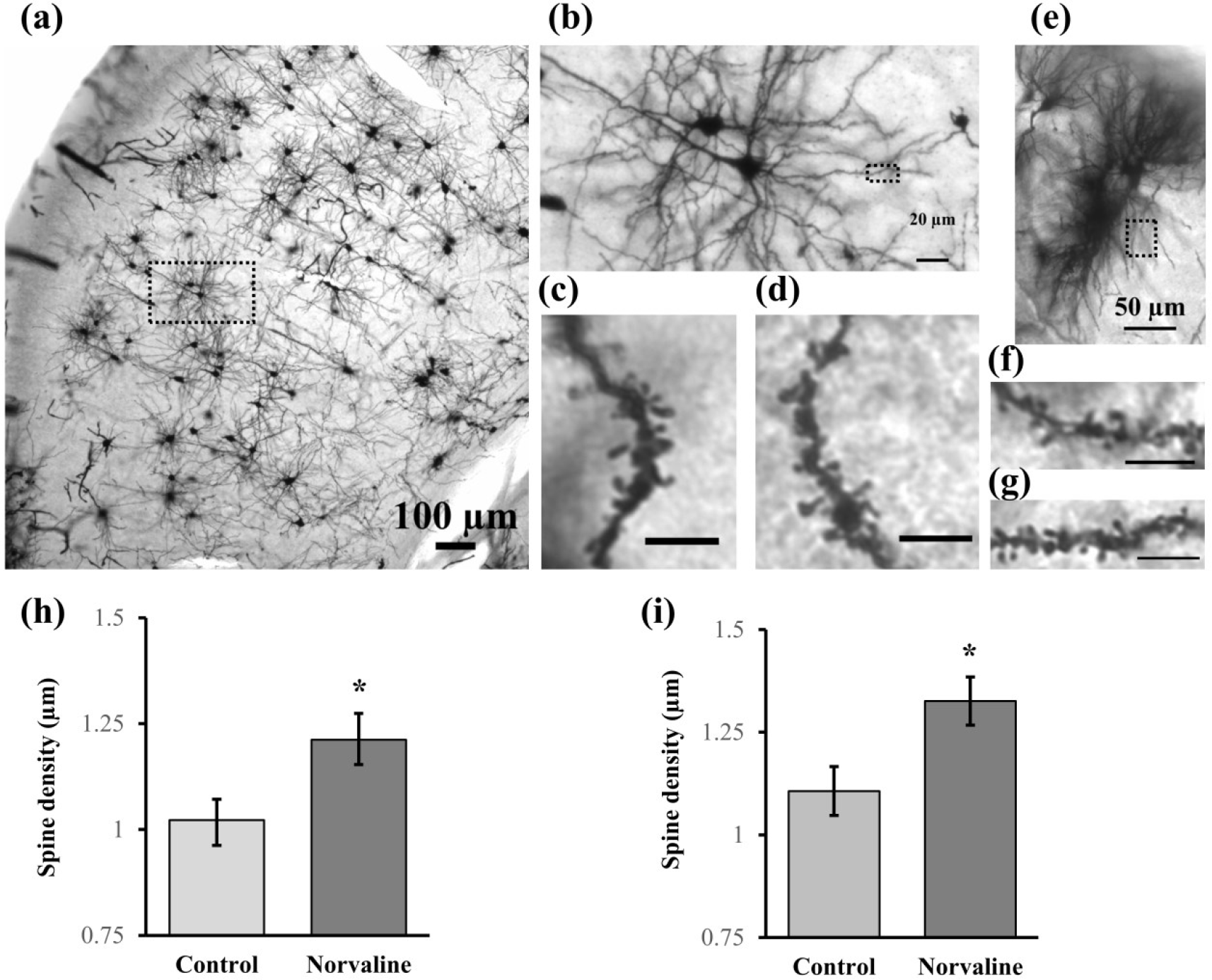
Photomicrographs of Golgi-stained 3xTg-AD mice cortical and hippocampal neurons from slices. (a) Representative Golgi-impregnated 10x image of the cortex. (b) Enlargement 20× of boxed area in (a) showing pyramidal neuron in the layer III. Representative image of the dendritic segments at 100× magnifications to display spines of pyramidal cells in control (c) and L-norvaline treated groups. Representative 20× image of the CA1 pyramidal cell. (f) Representative dendritic CA1 segments of control (f) and experimental group (g). Scale bar = 5 μm. Automated quantitation of spine density by Neurolucida software was subjected to statistical analysis by Student’s t-test. (h) Spine density of the cortical neuron.(i) Spine density of the hippocampal neuron. *p<0.05. The data are mean±SEM, n=36 for cortical neurons, n=24 for hippocampal cells (3 mice per group).

### L-norvaline amplifies expression rates of proteins related to synaptic plasticity

In order to gain more molecular insight into the behavioral phenotype, the effect of L-norvaline treatment upon the levels of neuroplasticity-associated proteins in 3xTg-AD mice was investigated with a set of proteomics assays. First, we tested the protein levels of 12 known important neuroplasticity related proteins in the hippocampus by western blot analysis (proteins listed in Supplementary Table 1). Second, an antibody array analysis was done to detect changes in protein levels at a whole-throughput level (approximately 1000 proteins). Western blot analysis revealed an increase of the synaptic and neuroplasticity related proteins in the L-norvaline treated group (Fig. 4a). Remarkably, expression of vesicular glutamate transporters 1 and 3 in the brain grew by 458 and 349% respectively, potentially increasing vesicular glutamate levels. Glutamate is the primary excitatory neurotransmitter and contributes to awareness and consciousness, which are essential for learning and forming memories. Protein microarray analysis revealed increased levels of other important neuron-related proteins following L-norvaline treatment. Notably, the expression of endothelial NOS increased significantly by 68%. Nitric oxide is a vasodilator and contributes to endothelial function and brain perfusion. Additionally, there was a substantial increase in cyclin E protein, which is a central component of the cell cycle machinery, and known to regulate synaptic plasticity and memory formation (Odajima et al. 2011).

**Figure 4.**
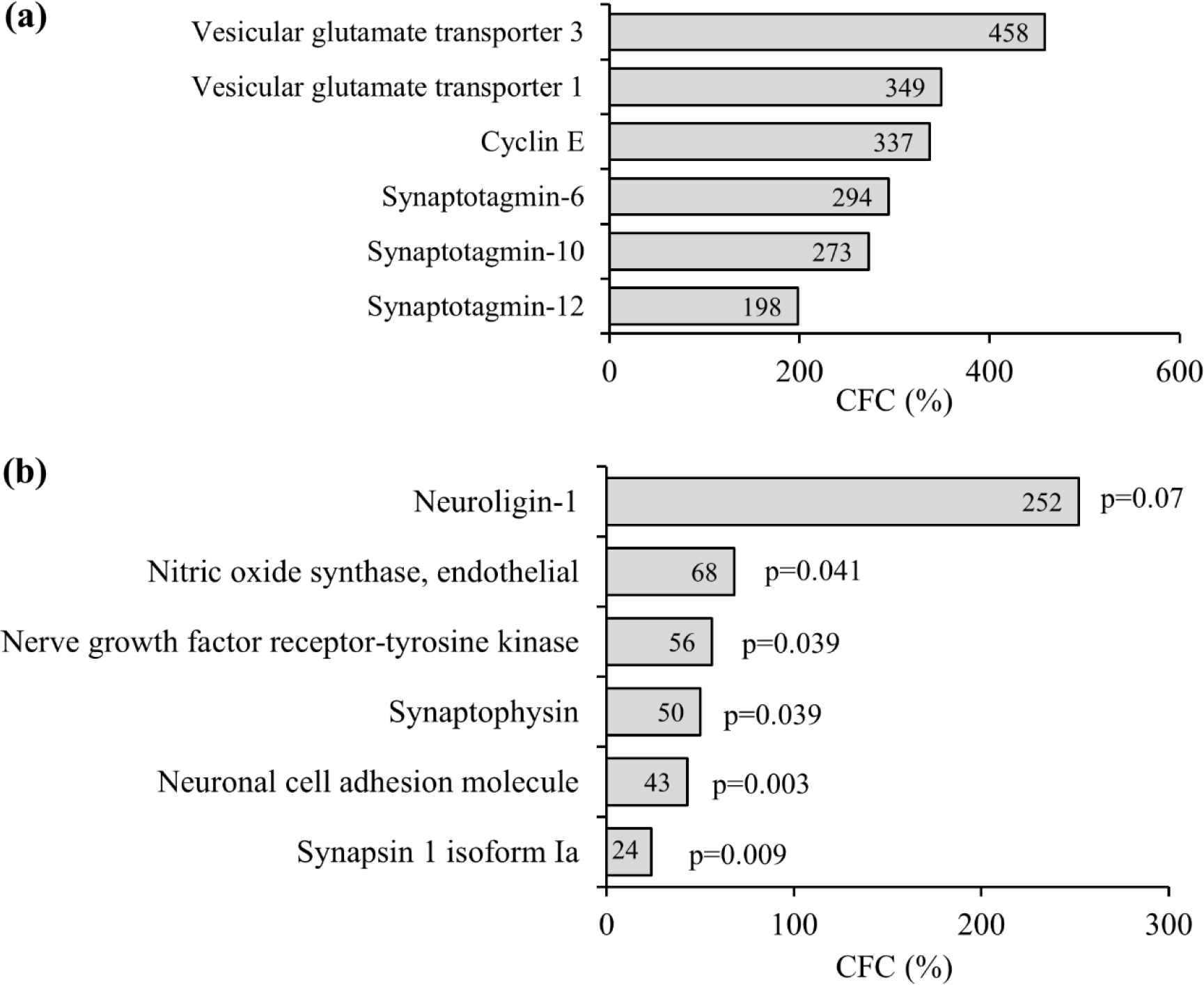
L-norvaline increases expression rate of neuroplasticity related proteins in the hippocampus of 3xTg-AD mice. Western blot summary with β-actin normalized trace quantities (a), antibody array selected data (b). CFC denotes change from control, p represents p-value of the change.

### L-norvaline reduces the rate of microgliosis in 3xTg-AD mice

Microglia act as the first and main form of active immune defense in the brain. An increase in microglia density is indicative of elevated pathogenic insults. Increased microglia density in the hippocampus of AD patients (Bachstetter et al. 2015) and 3xTg-AD mice was reported (Rodríguez et al. 2013). To assess the microglia density, image processing was performed using Zen Blue 2.5 (Zeiss) software with manually set threshold. First, the CA2-CA3 area with surface of 0.2 mm^2^ was subjected for the analysis. In order to guarantee selection of only the cells, which are entirely present in the acquired field, for IBA1 staining, cells with an area greater than 25 μm^2^ were considered for analysis, as widely accepted in the literature ((Zanier et al. 2015), (Zanier et al. 2016). A significant effect of the treatment upon “microglial density” and “Iba1+area fraction” was revealed by a One-Way ANOVA test (F(3,52)=4.64, p=0.006 and F(3,52)=4.098, p=0.01 respectively). Post-hoc Tukey’s Multiple Comparison Test indicated significant reduction (p<0.05) of the microglial density in the area of interest in the L-norvaline treated 3xTg-AD group (63.84±7.43 *versus* 83.66±5.11 cells/mm^2^, on average) (Fig. 5).

**Figure 5.**
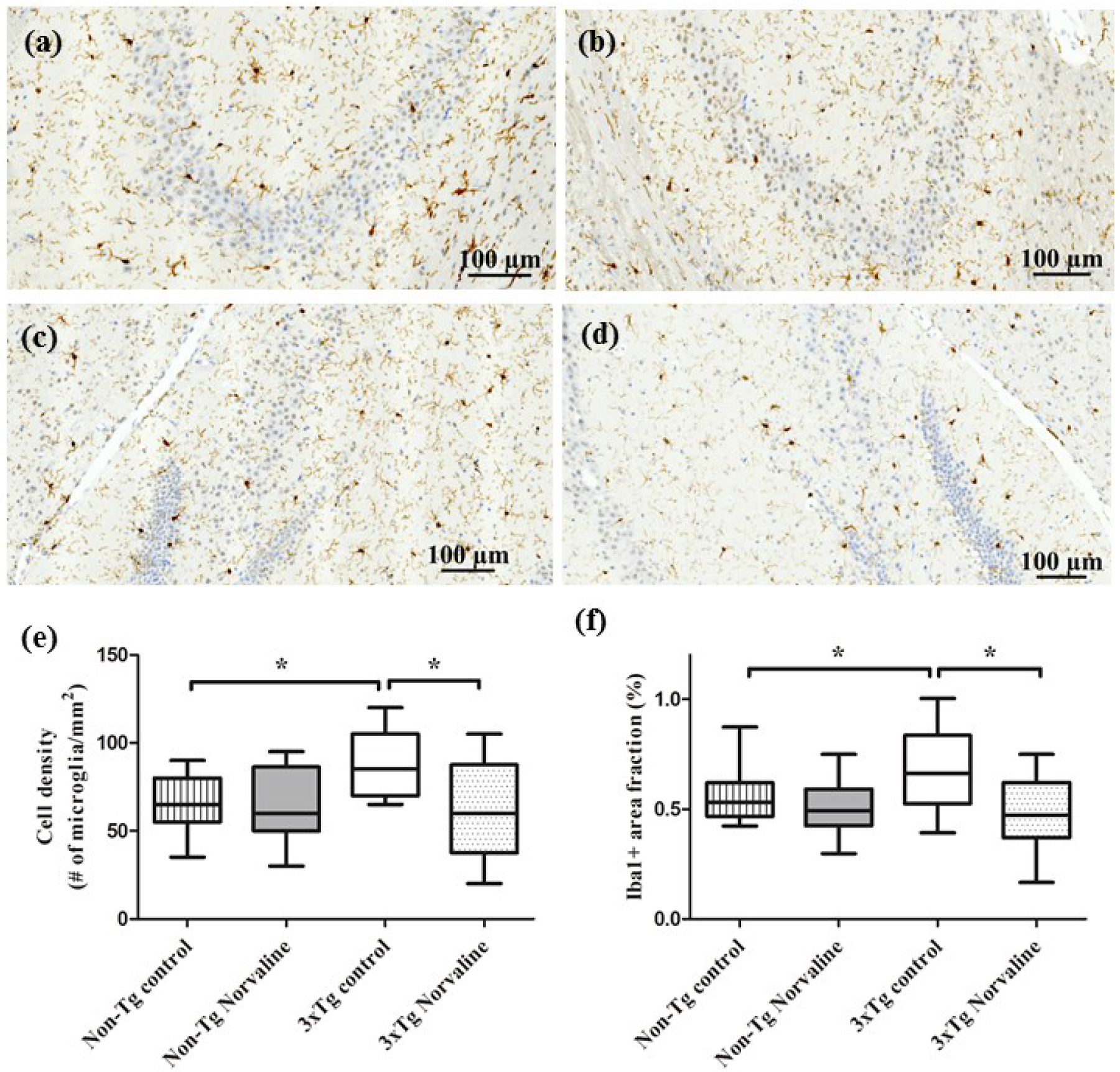
L-norvaline significantly decreases microglial density. Visualization and quantification of microglia with Iba1 staining in the hippocampus of 3xTg-AD mice (a, c). Representative hippocampal brightfield micrographs of CA1 and CA4 areas respectively of the control and (b, d) L-norvaline treaded 3xTg-AD mice (e) Box-plots showing the area density of microglia in the hippocampal CA3 of 3×Tg-AD mice and non-Tg control mice treated with a vesicle and L-norvaline. (f) Significant reduction in the area fraction (%) of IBA1 immuno-reactivity in the hippocampal CA3 of 3×Tg-AD mice treated with L-norvaline. *p<0.05 versus controls using a one-way ANOVA with Tukey’s post hoc test, n=20, four mice per group.

### Shape parameters of IBA1 positive cells situated in the hippocampus of treated with L-norvaline 3xTg mice indicate a prevalence of resting ramified cells compared to the untreated controls

Microglia morphology and function are closely related. Microglia are sensitive to the surrounding microenvironment and morphology is regulated by various stimuli (Fernández-Arjona et al. 2017). Neuroinflammation is characterized not only by increase in the microglial density, but by diverse morphological changes including de-ramification (E. J. Davis, Foster, and Thomas 1994). To address this issue several morphological features of the stained microglia were analyzed by using image analysis software (Image-Pro^®^ 10, Media Cybernetics, Inc. Rockville, USA) and ZEN 2.5. Iba1 is known to be expressed weakly by ramified microglia, however, upregulated and strongly expressed by activated microglia (Tynan et al. 2010), (Imai and Kohsaka 2002). So, the index of Integrated Optical Density (IOD) was measured to characterize Iba1 staining intensity (Schreiber et al. 2013). There was a significant effect of the treatment upon the microglial IOD (Fig. 6a). In the treaded group this parameter was reduced by 30%.

A more detailed analysis focusing upon the profile of microglial somata revealed several additional parameters that were significantly affected by the treatment. First, the average size (surface area) of the measured cells in the treated group contracted by 17% (50.92 ± 1.69 μm^2^ *versus* 60.93±1.06 μm^2^; p < 0.0001) (Fig. 6c). Second, the roundness or sphericity of the somata (4πArea/perimeter^2^), which is an aspect of microglial reactivity, was significantly increased in the treated with L-norvaline group (0.763 ± 0.012 *versus* 0.71 ± 0.005; p < 0.0037) (Fig. 6d). Additionally, the Feret’s diameter or max-min caliper was measured. The analysis reveals a significant diminution of Feret’s diameter in the treated group (6.82±0.182 μm *versus* 8.61±0.26 μm; p<0.0001) (Fig. 6b).

Distribution analyses furthermore revealed a left-shifted distribution of soma size and right-shifted distribution of circularity in the treated group (Fig. 6 e,f). Together, these findings suggest that the number of microglia with a small, round cell body (resting microglia) increases after the treatment. Cells in the control group have a bigger and more irregular soma shape (activated microglia). This observation accords with common classification of microglia morphology, which describes the resting cells as small, round cells with elaborate ramifications, and the activated microglia as bigger, more amoeboid cells with retracted processes (B. M. Davis et al. 2017).

**Figure 6.**
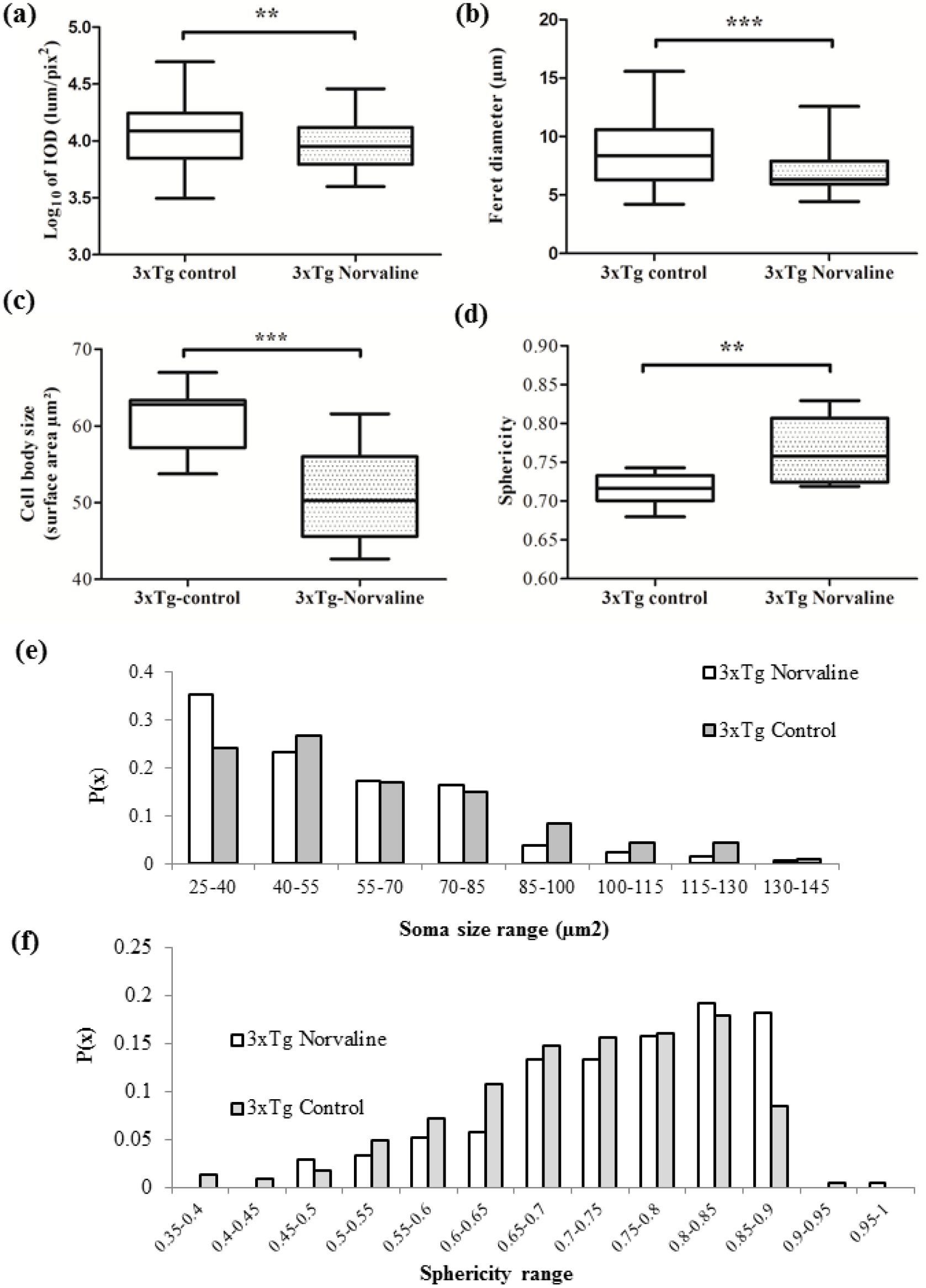
Quantitative characterization of microglia morphology in CA3 hippocampal area with S=0.2 mm2. (Five sections per mice were subjected for the analysis) (a) Integrated Optical Density, (b) Feret diameter, (c) Cell surface area, (d) Sphericity index, Frequency histogram of all microglial cells sampled in CA3 area with S=0.2 mm2. (e) Frequency distribution analysis of soma area shows a shift from large to smaller cell body sizes after the treatment; and (f) distribution analysis of circularity indicates that control cells are more likely to possess more irregular shape (i.e., to have a lower sphericity index).

### L-norvaline reverses astrocyte degeneration in brain areas with pronounced β-amyloidosis

Astrocytes were shown to modulate and control synaptic activity (Kimelberg and Nedergaard 2010). Glial fibrillary acidic protein (GFAP) is highly expressed in astrocytes (Jacque et al. 1978). We stained the brains and observed GFAP immunoreactive cells possessing typical stellate shape (Fig. 7e,f). We measured the astrocytes’ density within the hilus of 3xTg-AD mice by using ZEN 2.5. According to the literature, the average surface area of an astrocyte is about 250 μm^2^ (Williams, Grossman, and Edmunds 1980). To count the cells we filtered out all GFAP positive objects that less than 70 and more than 500 squared micrometers. There were no significant differences (p=0.62) observed in the density of GFAP+ astrocytes between the control and experimental group (Fig. 7h).

We quantified the levels of GFAP immunoreactivity (reflected by volume density) within different hippocampal areas. The comparative analysis with two-tailed Student’s t-test between experimental groups revealed significantly increased CA3 (p=0.0154) and CA4 (p=0.0008) glial somatic volumes in the animals treated with L-norvaline.

**Figure 7.**
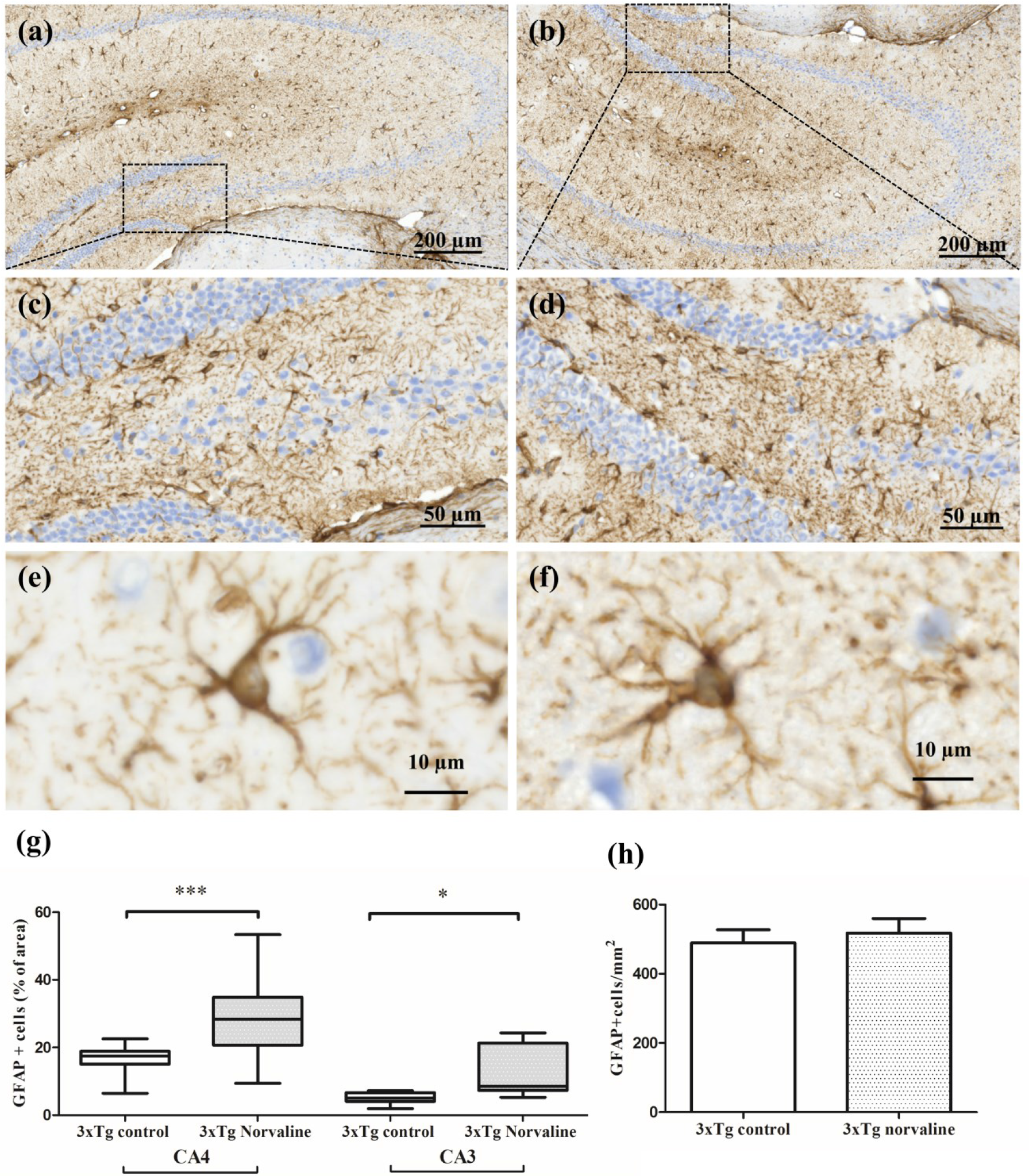
Treatment with L-norvaline does not lead to changes in GFAP-positive astroglial density in 3xTg mice; however, increases significantly the volume of astrocytes’ somata. Representative hippocampal brightfield 20x micrographs of control (a) and L-norvaline treated mice. CA4 area of control (c) and treated (d) 3xTg-AD mice. Representative 40x images of single astrocyte from CA4 area of control (e) and treated (f) mice. (g) Significant reduction in the area fraction (%) of GFAP+ cells in the hippocampal CA3, CA4 areas of control 3×Tg-AD mice (g) and treated with L-norvaline (h). versus controls (h) GFAP+ cells density in hilus. n=20, four mice per group. *p<0.05, **p<0.01, ***p<0.001.

### ARGI is distinctly expressed in the areas with pronounced amyloidosis. L-norvaline treatment effectively reduces ARGI expression

We examined ARGI protein spatial expression within the hippocampi of 3xTg-AD mice, and its spatial relationship with Aβ deposition using immunocytochemistry (Fig. 8). ARG1 is preferentially accumulated in the CA1-CA4 areas of the hippocampus and primarily detected inside the cells (Fig 7g). Treatment with L-norvaline led to a significant (p=0.0073) reduction in the overall ARG1+ cell surface area and IOD (p=0.0004), which reflect the decrease in the levels of ARG1 protein in the brains of the treated animals.

**Figure 8.**
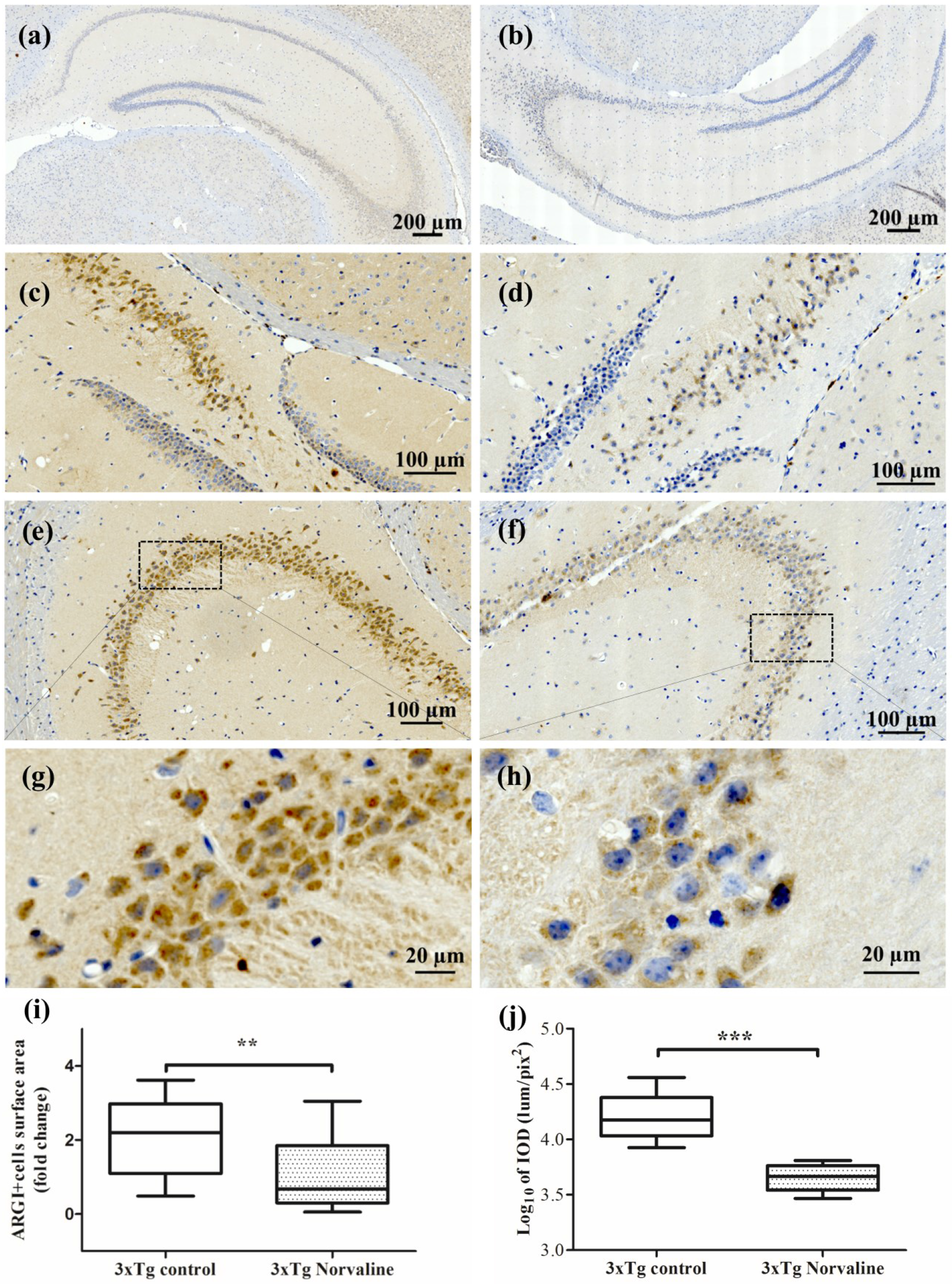
L-norvaline significantly decreases ARGI immunopositivity. Representative hippocampal bright-field 20x micrographs of control (a) and L-norvaline treated mice(b). CA4 area of control (c) and treated (d) 3xTg-AD mice. CA3 area of control (e) and treated (f) mice with 40 x insets (g) and (h) respectively. (i) ARGI+ cells surface area in CA3 (s=0.2 mm^2^). (j) Integrated optical density of ARGI+ cells in CA3. (n=24, with 4 mice per group)

### Mitochondrial ARGII is expressed in the cytosol of the of CA2-CA3 hippocampal cells

We detected augmented ARGII immune-positivity of the cells situated in the CA2 hippocampal area of 3xTg mice. This pattern of the staining was observed in the treated and control group as well. Mitochondria were seen with 40x and more magnification (most clearly in the dentate gyrus) without significant differences in numbers and intensity between the groups. We did not detect significant effect of the treatment upon “ARGII+ objects surface area”, however the index of IOD was reduced significantly (p<0.05) in the treated with L-norvaline mice.

**Figure 9.**
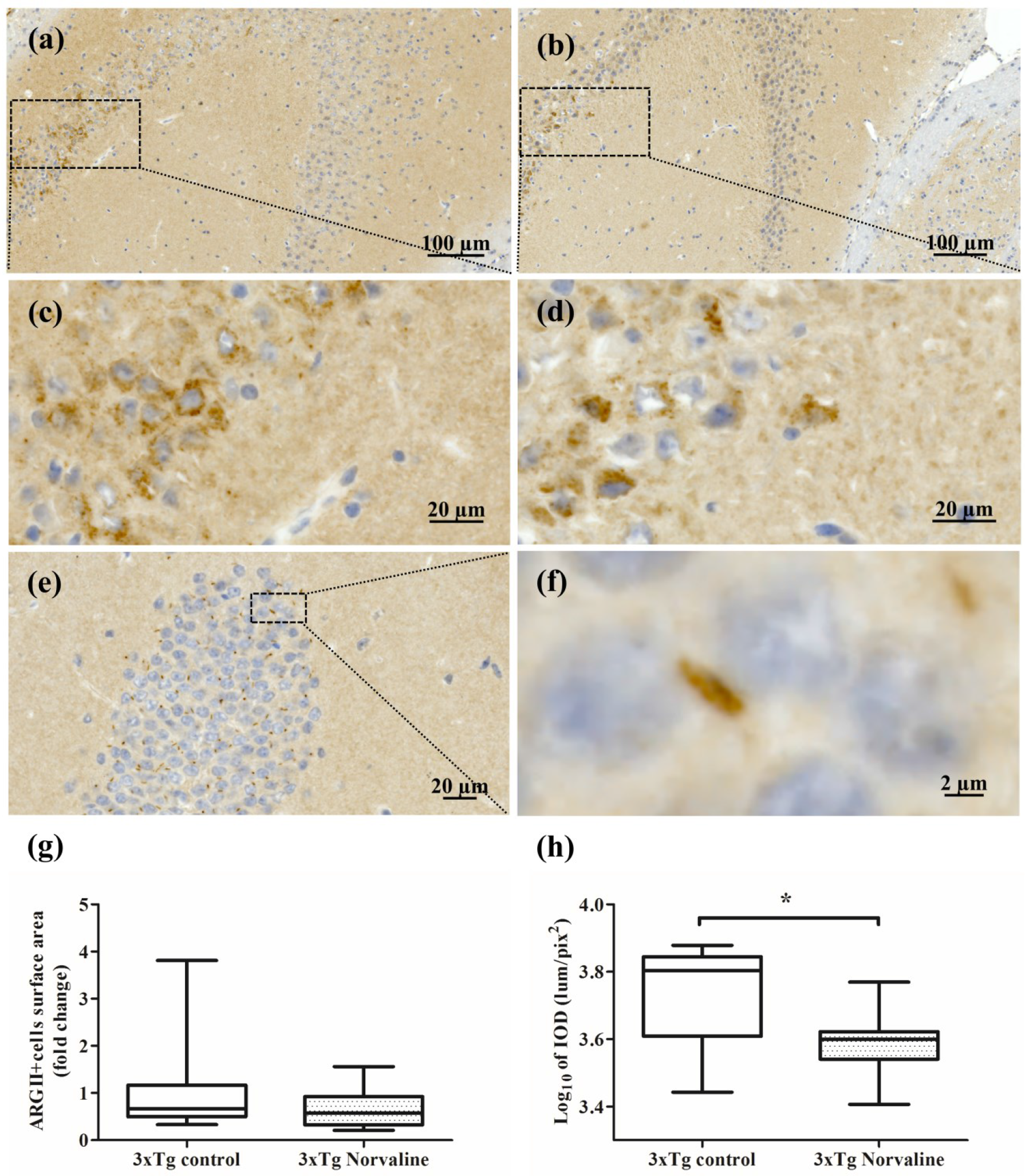
Representative hippocampal CA2-CA3 bright-field 20x micrographs of control (a) and L-norvaline treated mice (b). CA2 40x insets of control (c) and treated (d) mice. (e) Dentate gyrus representative 20x image with mitochondria stained with ARGII antibody (f) 40x zoomed in inset with an evidently seen mitochondrion. (g) ARGII+ cells surface area in CA2 (s=0.2 mm2). (h) Integrated optical density of ARGII+ cells in CA2. (*p<0.05, n=12, with 4 mice per group).

## Discussion

In this study, we treated 3xTg-AD mice with a non-proteinogenic unbranched-chain amino acid L-Norvaline. This substance possesses arginase-inhibiting properties (Pokrovskiy et al. 2011). Our results indicate that L-norvaline is well tolerated by mice and a long-term treatment with the substance does not lead to detectable behavioral changes or loss of weight in wild type animals. We show that 3xTg-AD mice from the L-norvaline treated group not only exhibit improved spatial learning and memory but also demonstrate reduction of cerebral Aβ burden together with substantial decrease of Aβ toxic fibrillary and oligomeric species. Moreover, we report a decrease in microglial density in treated 3xTg-AD mice with a shift from activated to resting phenotype of the microglia.

The idea to block L-arginine depletion and to reverse its deprivation, thereby halting memory loss, and reducing AD symptoms has previously been explored. For example, Kan *et al.* used DFMO, a relatively toxic, mostly irreversible, inhibitor of ornithine decarboxylase. DFMO was provided together with putrescine in a CVN mouse model of AD (Kan et al. 2015). In our approach, we inhibit arginase directly by using L-norvaline, which is structurally similar to ornithine (a product of arginase) and acts via a negative feedback inhibition mechanism. Accordingly, we did not administer the animals with polyamines to avoid the undesirable side effects of DFMO and provided the mice with norvaline dissolved in water in their home cages, which is a much less invasive methodology compared to forced gavage feeding. Furthermore, we utilized the universally recognized animal model of AD.

The mice from L-norvaline treated group showed a very strong phenotype in two different paradigms, which supports the sound effect of the treatment on short and long-term spatial memory acquisition. Moreover, we disclose a significant reduction in the rate of 6E10 positivity in the cortex of 3xTg mice treated with L-norvaline. Additionally, we subjected hippocampal tissue homogenates to western blot analysis to determine the composition of the soluble Aβ fraction. We evidence that the hippocampus of the six-seven-month-old 3xTg mice treated with L-norvaline harbors significantly lower levels of oligomeric and fibrillary conformers, compared to untreated 3xTg tissue homogenates. Growing evidence indicates that soluble low weight Aβ oligomers and intermediate aggregates are the most neurotoxic forms of Aβ triggering synapse dysfunction and loss, and neuronal damage, which manifest consequentially in behavioral attributes of AD such as memory deficits (Walsh and Selkoe 2007). Our findings accord with these phenomena.

We observed a significantly increased ARGI expression in the areas with pronounced Aβ deposition. There is a consensus that the expression of two isoforms of arginase can be compelled by various stimuli (Sidney and Morris 1992), (Lange et al. 2004). In the brain of the AD model mice ARGI was shown to be localized not only in the cells, but to be distributed in the extracellular space. In the hippocampus, it displays a spatial correlation with Aβ deposition, and Iba-1 expression (Kan et al. 2015). ARGII is inducible by a variety of factors as well, including LPS, TNFα, oxidized LDL, and hypoxia (Ryoo et al. 2011). Moreover, it was confirmed that its activation is associated with translocation from the mitochondria to the cytosol (Pandey et al. 2014).

We applied a set of immunohistochemistry (IHC) approaches to display significantly increased ARGI levels in the CA2-CA4 hippocampal areas of the 3xTg-AD mice, where we detected the most distinct intracellular Aβ deposition. These findings accord with the current literature (Kan et al. 2015). Of note, in our case, ARGI staining was primarily localized inside the cells, which unlike the findings presented in the cited above paper, could be attributed to differences in used murine model of AD. Furthermore, we show that L-norvaline treatment significantly reduces the rate of ARGI expression in the hippocampi.

Additionally, we observed augmented ARGII immunopositivity in the cells of CA2 area of 3xTg mice, which indicates its translocation from the mitochondria to the cytoplasm, supposedly by the same mechanism that was shown previously (Pandey et al. 2014). Still, we did not detect differences between the groups in the relative surface area of ARGII positive objects located in CA2, however, optical density of the objects was reduced significantly in the treated mice.

Chronic neuroinflammation is a prominent feature of AD pathogenesis (Heneka et al. 2015). Microglial proliferation is increased in AD patients as well as AD animal models and positively correlates with disease severity (Clayton, Van Enoo, and Ikezu 2017). Likewise, it was suggested that inflammation correlates better with cognitive decline than accumulation of Aβ (Holmes et al. 2009). Consequently, there have been attempts to interfere with inflammatory complement-mediated processes leading to synaptic loss in AD mice. Spangenberg *et al.* showed that eliminating microglia in AD mice prevents neuronal loss without modulating Aβ pathology (Spangenberg et al. 2016). Another study utilizing pharmacological targeting of the colony-stimulating factor 1 receptor demonstrated that inhibition of microglial proliferation prevents the progression of AD pathology (Olmos-Alonso et al. 2016). These results led to the hypothesis that depleting microglia and blocking complement pathway activation are beneficial for cognitive functions.

In our study, we observed a more than 30 % increase of Iba1 immunopositive cells density in the CA3 area of the seven-month-old 3xTg mice hippocampi compared to the age and sex-matched non-Tg controls. Moreover, we confirm that cognitive function improvement in the treated with L-norvaline group is associated with significant reduction of the hippocampal microgliosis.

There is a strongly grounded hypothesis that activated microglia cause synaptic and wiring dysfunction by pruning synaptic connections (Hong, Dissing-Olesen, and Stevens 2016), so it was proposed to target the microglia-synapse pathways to accomplish an early prevention of AD symptoms (Xie et al. 2017). We report a substantial increase in dendritic spine density in the cortices and hippocampi of the 3xTg mice treated with L-norvaline. Moreover, we evidence a significant escalation of the pre- and postsynaptic proteins’ levels in the experimental group, which we associate with the increase in spine density.

In parallel with microgliosis and astrogliosis (Osborn et al. 2016), the development of AD is associated with astrodegeneration (Rodriguez et al. 2009). Astrocytes are essential for neurotransmission. They provide neurons with glutamine, which is a precursor molecule for glutamate and GABA (Rose, Verkhratsky, and Parpura 2013). Consequently, astrodegeneration leads to synaptic strength deficits and contributes to a decline in the number of active synapses, observed at the early stages of AD (Verkhratsky, Zorec, Rodriguez, et al. 2016). Morphological astrocytes’ atrophy has been noted in the hippocampi and cortices of the 3xTg-AD mice (Yeh et al. 2011). This atrophy is characterized by reduction of cell volume, and a decrease in the number of processes (Verkhratsky, Zorec, Rodríguez, et al. 2016). Interestingly, the density of astrocytes remains relatively stable in the hippocampus, and cortices of the 3xTg mice for more than one year and is comparable to the non-Tg controls (Kulijewicz-Nawrot et al. 2012). It was reported by Olabarria *et al.* that six-month-old 3xTg-AD mice demonstrate apparent reduction of the astrocyte somatic volume in DG, which becomes significant at 12 months (Olabarria et al. 2010). Several *in vitro* studies evidenced that Aβ is capable of triggering astroglial Ca^2+^ signaling, which leads to mitochondrial depolarization, and oxidative stress (Andrey Y Abramov, Canevari, and Duchen 2003) (A. Y. Abramov 2004).

Our results accord with the actual data showing astrodegeneration in the areas with intense Aβ deposition. Moreover, we demonstrate that treatment with L-norvaline leads to a significant 75% increase in the GFAP-positive surface area in the CA3 region. We speculate that the observed reduction of the astrocyte somatic volume is dependent upon the emergent Aβ deposition and subsequent local oxidative stress. We further hypothesize that the detected increase of astrocyte somata volume in the experimental group is associated with: reduction in the concentration of toxic oligomeric and fibrillary Aβ species, direct cytoprotective effect of L-norvaline, changes in L-arginine bioavailability.

L-norvaline possesses multiple mechanisms of activity. Ming *et al.* elegantly demonstrated that L-norvaline inhibits TNFα-induced endothelial inflammation, independently of NOS and arginase activity. The substance reduces the expression of the inflammatory adhesion molecules (VCAM-1, ICAM-1, E-selectin) in the presence of TNFα *in vitro* (Ming et al. 2009). The authors utilized RNA interference technology to demonstrate that the anti-inflammatory properties of L-norvaline are attributed to its ability to inhibit S6K1 kinase, which is involved in the regulation of protein synthesis. Subsequently, Caccomo *et al.* evidenced that S6K1 activity is higher in 3xTg-AD mice compared to WT (Caccamo et al. 2015). Moreover, removing one copy of the S6K1 gene restores S6K1 activity in 3xTg-AD mice to the non-Tg levels. Additionally, the levels of phosphorylated S6K1 are higher in 3xTg-AD mice compared to their WT. Caccomo *et al.* also showed that removing one copy of the S6K1 gene from the 3xTg-AD mice is sufficient to rescue the LTP deficits. They speculate that LTP improvement corresponds to the changes in synaptic marker synaptophysin, which is reduced in 3xTg-AD mice and escalates dramatically in S6K1 deficient mice.

In our study, we demonstrate a significant (p=0.039) elevation by 50% of the levels of synaptophysin in the hippocampus of the animals treated with L-norvaline that coheres with the mentioned above data. Therefore, we verify that S6K1 is a key protein partially responsible for hippocampal synaptic deficit found in 3xTg-AD mice and its inhibition by L-norvaline is sufficient to rescue spatial memory deficits in this model of AD. Noteworthy observation of a significant (p=0.038) reduction by 53% of RAC-alpha protein-serine/threonine kinase (Akt1) levels in the treated group (Supplementary Table 2). This kinase is a key modulator of the AKT-mTOR signaling pathway and regulates many processes including metabolism, proliferation, and cell survival (Memmott and Dennis 2009). It is located upstream to mTOR and S6K1, thus downregulation of Akt1 can negatively influence S6K1 activity. Upregulation of mTOR-signaling pathway is known to play a central role in major pathological processes of AD (Wang et al. 2014). Consequently, mTOR-signaling inhibition is a novel therapeutic target for AD (Z. Cai et al. 2015) (Tramutola, Lanzillotta, and Di Domenico 2017). There is still much work to be done in order to decipher the precise mechanisms of L-norvaline regulation of the AKT-mTOR signaling pathway, which is beyond the scope of this paper.

Caccomo *et al.* demonstrate that S6K1 deficiency lowers Aβ pathology in 3xTg-AD mice (Caccamo et al. 2015). They disclose particularly, that levels of C terminal fragment 99 (C99) of APP were significantly lower in 3xTg-AD/S6K1^+/−^ mice compared to the regular 3xTg-AD mice. Previously, C99 was identified by Lauritzen *et al.* as the earliest βAPP catabolite and main contributor to the intracellular βAPP-related immunoreactivity in 3xTg mice, which is implicated in the neurodegenerative process and cognitive impairment taking place in this animal model of AD (Lauritzen et al. 2012). In order to explain the phenomenon, Caccomo *et al.* measured BACE-1 protein levels using Western blot and BACE-1 mRNA by qPCR. The authors concluded that low S6K1 signaling reduces BACE-1 translation and consequently decreases generation of the C99 fragment. This phenomenon partially explains our results showing significantly reduced levels of toxic Aβ species in the brains of the treated with L-norvaline mice.

We found that superoxide dismutase [Cu-Zn] levels were elevated by 19% in the brains of the treated animals (Supplementary Table 2). This enzyme plays a critical role in cellular response to reactive oxygen compounds. It was convincingly demonstrated in a rat model with NMDA excitotoxic lesion that overexpression of superoxide dismutase [Cu-Zn] leads to neuroprotection and improved functional outcome (Peluffo et al. 2006). Moreover, we observed significantly elevated by 24% the expression rate of phosphatidylinositol 3-kinase regulatory subunit alpha. Yu *et al.* verified that this kinase plays a role in protection against H_2_O_2_-induced neuron degeneration (X. Yu et al. 2004). The data suggest that L-norvaline activates the cellular protection mechanisms against oxidative stress.

Overall, our results provide compelling evidence indicating that L-norvaline is a multifunctional agent and a potential drug for the treatment of AD. It appears to be a promising candidate for clinical development. The functional effects of the substance are multifarious and there is much work to be done to disclose all of its potentials and the precise mechanisms of action.

## Materials and methods

### Strains of mice and treatment

The homozygous 3xTg-AD mice harboring a mutant APP (KM670/671NL), which is a human mutant PS1 (M146V) knock-in, and tau (P301L) transgenes (B6;129-Psen1^tm1Mpm^ Tg(APPSwe, tauP301L)1Lfa/J) were obtained from Jackson Laboratory^®^ and bred in our animal facility. The mice exhibit synaptic deficiency with both plaque and tangle pathology (Oddo et al. 2003). Four-month-old homozygous 3xTg-AD mice and age matched male C57BL/6 mice were used as non-transgenic controls (Non-Tg) in all experiments. C57BL/6 mice are wildly accepted as the non-transgenic controls of 3xTg-AD mice (Lin et al. 2018), (Lee et al. 2015), (Janelsins et al. 2005), (Baek et al. 2016). Randomly chosen animals were divided into four groups (14-15 mice each) and provided with water and food *ad libitum*. L-norvaline (Sigma) was dissolved in water (250 mg/L) and supplied in the home cages. Taking into account that average consumption of water by mice in laboratory conditions equals 5-6 ml/day (Bachmanov et al. 2002), for the 3xTg-AD mice with average weight of 33 grams, the dose was about 1.5 mg/day. The mice underwent appropriate treatment and were behaviorally tested (Supplementary Fig. 1). We measured the weight of the animals every week during the experiment and did not observe any significant effect of the treatment upon the weight. All animal housing and procedures were performed in compliance with instructions established by the Israeli Ministry of Health’s Council for Experimentation on Animals and with Bar Ilan University guidelines for the use and care of laboratory animals in research. The experimental protocol was approved by the Committee on the Ethics of Animal Experiments of the Bar Ilan University (Permit Number: 82-10–2017).

### Behavioral tests

#### Morris water maze (MWM)

The MWM task was conducted to measure long-term learning and memory function as described previously (Buccafusco 2009). Briefly, a black plastic pool with a diameter of 120 cm and a height of 50 cm was filled with water (24 ± 2°C) and rendered opaque by adding skim milk powder (Sigma). A cylindrical dark colored platform with a diameter of 10 cm was placed 0.3 cm under the water surface and kept consistently within one of the quadrants. Training was conducted for seven consecutive days with two trials a day.

The test consists of three separate procedures (Bromley-Brits, Deng, and Song 2011). On the first day, the animals were tested with visible platform. A flag was placed on the platform to increase its visibility (Bromley-Brits, Deng, and Song 2011). Visible platform test identifies gross visual deficits caused by the treatment that might confuse interpretation of the results obtained from the MWM experiment (Buccafusco 2009). The mice underwent two trials with 90 sec cut-off each and a 15 minutes interval between trials. On days 2-6, the animals were tested with hidden under water platform (without a flag). Each animal was placed into the pool at one of four locations (poles). The mice were allowed to find the platform within 90 seconds and stay on it for 15 seconds. If a mouse could not escape during the time limit, it was directed gently towards the platform. The probe test without the platform was performed on the seventh day. The mice were released in the center of the maze and subjected to a single trial with free swimming during 90 seconds.

#### Spontaneous alternation Y-Maze

Short-term working memory was assessed in the Y-maze consisted of three arms (each of 35 cm long, 25 cm high and 10 cm wide) at a 120° angle from each other. After introduction to the center of the maze, each animal was allowed to freely explore the maze for 8 min. The maze was cleaned with a 10% aqueous ethanol solution between each trial. Spontaneous alternation was defined as a successive entry into three different arms, on overlapping triplet sets. An arm entry was counted when the hind paws of the mouse were completely within the arm (Knight et al. 2014).

#### Video tracking

All behavioral experiments were recorded by the Panasonic WV-CL930 camera with the Ganz IR 50/50 Infrared panel. The recorded video files were analyzed with the Ethovision XT 10 software (Noldus^®^) by a blind to the treatment schedule experimenter.

#### Tissue sampling

After the behavioral tests 3xTg mice were deeply anesthetized with sodium pentobarbital (60 mg/kg), administered intraperitoneally and decapitated. Their brains (four from each group) were sliced (0.5 mm thick) immediately in the mouse brain slicer matrix. Sections (between 1.7 mm and 2.2 mm posterior to bregma according to the atlas of Franklin and Paxinos) were used for sampling (Franklin and Paxinos 2008). The hippocampi were punched in the region of dentate gyrus using a13-gauge microdissection needle on each hemisphere, frozen and stored at −80°C.

#### Antibody microarray

The assessment of “hit” proteins’ expression was performed by the use of the Kinex KAM-1150E Antibody Microarray (Kinexus Bioinformatics), in accordance with the manufacturer specification. The analyses were done with hippocampal lysates as described on Kinexus’ web page (www.kinexus.ca). Briefly, lysate protein from each sample (100 μg) was labeled covalently with a fluorescent dye combination. Free dye molecules then were removed via gel filtration. After blocking non-specific binding sites on the array, an incubation chamber was mounted onto the microarray to permit the loading of the samples. After the incubation, unbound proteins were washed away. Two 16-bit images have been captured by a ScanArray Reader (Perkin-Elmer) for each array.

#### Golgi stain procedure

For Golgi staining, three 3xTg-AD mice per group were perfused transcardially with 0.1 M phosphate-buffered saline (pH 7.4) and brains were processed using superGolgi Kit according to the manufacturer’s protocol (Bioenno Lifesciences, Santa Ana, CA, USA). Briefly, brains were immersed in impregnation solution for 11 days, followed by 2 days incubation in a post-impregnation solution. Once the impregnation of neurons was complete, the brain samples were coronally sliced (150 μm) using a vibrating microtome (Campden Instruments) and serially collected in a mounting buffer. Then they were mounted upon 1% gelatin-coated glass slides and stained with the staining solution and coverslipped using Permount (Fisher Scientific, Houston, TX). Two corresponding sections (between 1.7 mm and 2.0 mm posterior to bregma according to the atlas of Franklin and Paxinos), representing the barrel cortex and the hippocampus, per animal were chosen for stereological analysis (Franklin and Paxinos 2008).

Upright Microscope ApoTome (Quorum Technologies) was used for imaging. To assess dendritic morphology, low magnification (10x/0.3, 20 x/0.8, and 40x/0.75 lens) images (Z-stack with 0.5 μm intervals) of pyramidal neurons, with cell bodies located in the cortical layer III and the CA1 region of the hippocampus, were captured by ORCA-Flash4.0 V3 camera. To evaluate dendritic spine morphology, high magnification (100x/1.4 oil objective with digital zoom 3) images (Z-stack with 0.25 μm intervals) were taken.

#### Spine density measurement

Captured images were coded and a blind to the experimental conditions investigator performed the analysis. The quantification was accomplished by use of Zen Blue 2.5 (Zeiss) and Neurolucida 360 version 2018.1.1 (MBF Bioscience, Williston, Vermont, USA). The software allows an automatic unbiased quantitative 3D analysis of identified neurons and was utilized to determine the number of spines per dendritic length (Dickstein et al. 2016). Spine detection threshold was set with outer range of 2.5 μm, minimal height of 0.3 μm, sensitivity of 100%, and minimal count of 10 voxels.

For each mice hemisphere, three cortical and two hippocampal pyramidal cells with somata located in the center of the 150-μm corresponding sections were selected for analysis (36 and 24 neurons per experimental group respectively). Spine density was quantified on secondary basal dendritic segments located farther than 40 μm from the soma of the layer III pyramidal cells and secondary oblique dendritic branch localized in the stratum radiatum of the CA1 hippocampal pyramidal neurons.

#### Immunostaining

Seven animals from each group were deeply anesthetized and transcardially perfused with 30 ml of phosphate buffer saline (PBS), followed by 50 ml of chilled paraformaldehyde (PFA) 4% in PBS. The brains were carefully removed and fixated in 4% PFA for 24 hours. Then, the brain from three 3xTg mice per group were transferred to 30% sucrose in PBS at 4°C for 24 hours and frozen at −80°C. The brains from other four mice (including non-Tg) per group were dehydrated and paraffin-embedded.

### Beta-Amyloid, 1-16 (6E10) Antibody Staining

The brains were sliced on a Leica CM3050 S cryostat to produce 30 μm floating sections. The corresponding sections (between 1.7 mm and 2.2 mm posterior to bregma) were chosen for analysis. These sections were blocked and incubated overnight with primary purified anti-β-Amyloid, 1-16 Antibody 6E10 (1:100, Biolegend), at 4°C, followed by washing and incubation with secondary antibodies Alexa555 goat anti-mouse IgG (1:200, Invitrogen) at room temperature during 1 hour, and Hoechst (1:5000, Sigma) for 3 min.

### Quantitative histochemical analysis of microglia (Iba1), astrocytes (GFAP); ARGI & ARGII

The paraffin-embedded tissue blocks were chilled on ice and sliced on Leica RM2235 manual rotary microtome at a thickness of 4 μm. The sections were mounted onto gelatin-coated slides, dried overnight at room temperature, and stored at 4 °C in slide storage boxes.

For quantitative histochemical analysis of microglia (Iba1), astrocytes (GFAP), ARGI & ARGII immunopositivity five brain coronal sections per mouse, per staining (at 1.9-2.0 mm posterior to bregma) were used. Staining was performed on fully automated Leica Bond III system (Leica Biosystems Newcastle Ltd, UK). Tissues were pretreated with epitope-retrieval solutions (ER, Leica Biosystems Newcastle Ltd, UK) followed by 30 minutes incubation with primary antibodies. Iba1 antibody (Novus biologicals, #NB100-1028) was diluted 1:500, GFAP (Biolegend, #835301) – 1:1000, ARGI (GeneTex, GTX113131) – 1:500, and ARGII (GeneTex, #GTX104036) – 1:600. The Leica Refine-HRP kit (Leica Biosystems Newcastle Ltd, UK) used for detection and counter-stain with Hematoxylin. Omission of Iba1, GFAP, ARGI, and ARGII primary antibodies served as negative control. For positive control of ARGI we stained the mouse liver tissue and kidney for ARGII.

### Imaging and quantification

Sections were mounted and viewed under the slide scanner for fluorescence and bright field Axio Scan.Z1 (Zeiss) with 40x/0.95 objective. Images were taken with Z-stack of 0.5 μm. Additionally, Axio Imager 2 Upright ApoTome Microscope was used to capture images with 100x/1.4 oil immersion objective. Immunolabeling was examined in the corresponding hippocampal areas, and image analysis was performed with Zen Blue 2.5 (Zeiss) and Image-Pro^®^ 10 (Media Cybernetics) software with fixed background intensity threshold for all sections representing a single type of staining.

### Morphometric cell analysis

The resting microglia exhibit a ramified phenotype, and adopt different morphologies, ranging from a highly ramified to an amoeboid-like phenotype upon activation (Kozlowski and Weimer 2012), (Torres-Platas et al. 2014). To quantify these morphological changes we defined a list of morphological parameters, which are accepted in the literature and expected to capture the shift (Zanier et al. 2015). These include cell perimeter length, Feret diameter, soma size, and sphericity ( 4π×area/ perimeter ^2^). To assess the staining density we measured integrated optical density (IOD) computed as mean density per pixel area (lum/pix^2^) (Carleton et al. 2018). IOD values of Iba1 stained microglia from CA3 regions with S=0.1 mm2 were calculated as log_10_ of the values acquired with Image-pro^®^ 10 (Carleton et al. 2018). Two appropriate sections (bregma −1.8 and −2.0 mm) per mice were subjected for the analyses with total amount of 125 cells for untreated and 90 cells for L-norvaline treated groups.

### Western blotting

To study the rate of beta-amyloidosis we examined the A11 and OC immunoreactivity of the hippocampal lysates. A11 is an antibody that detects the conformation of amyloid oligomers irrespective to their amino acid sequence (Kayed et al. 2003) and anti-amyloid fibrils OC antibody recognizes fibrillary forms.

Additionally, we analyzed and validated the relative difference in the expression levels of various neuroplasticity related proteins that revealed significant changes in the Antibody microarray assay. The set-up and the antibody nomenclature are presented in the supplementary Table 2.

Protein concentration was determined using the Bradford assay. Fifteen μg of pooled from four brains (each group) hippocampal lysates were analyzed by the Kinetworks^TM^ Custom Multi-Antibody Screen KCSS-1.0 as described by the manufacturer. Briefly, the Kinetworks^TM^ analysis involves resolution of a single lysate sample by SDS-PAGE and subsequent immunoblotting with validated antibodies. Antibodies bound to their target antigen on the nitrocellulose membrane are detected using the ECL Detection System.

### Statistical analysis

Statistical analyses were conducted using SPSS (version 22, IBM, Armonk, NY) for Windows. The significance was set at 95% of confidence. All results are presented as mean with standard error. Data were shown to fit a normal distribution using the Shapiro-Wilk test for normality, and Levene’s test was used to confirm equal variance between groups being compared.

For the Y-maze, visible platform test and probe trials (MWM) the means are compared between groups via a one-way ANOVA with the Tukey’s Multiple Comparison test used for post-hoc analyses. For the hidden platform test, the escape latencies of two trials were averaged for each mouse each day and then analyzed across the five days of testing (Buccafusco 2009). A one-way repeated-measures analysis of variance (ANOVA) was applied with day as the repeated measure and latency as the dependent variable. For comparison between groups, Tukey’s Multiple Comparison test was used when main analysis revealed significance. Student’s t-test was used to compare means between two groups.

## Supplemental material

**Figure S1.**
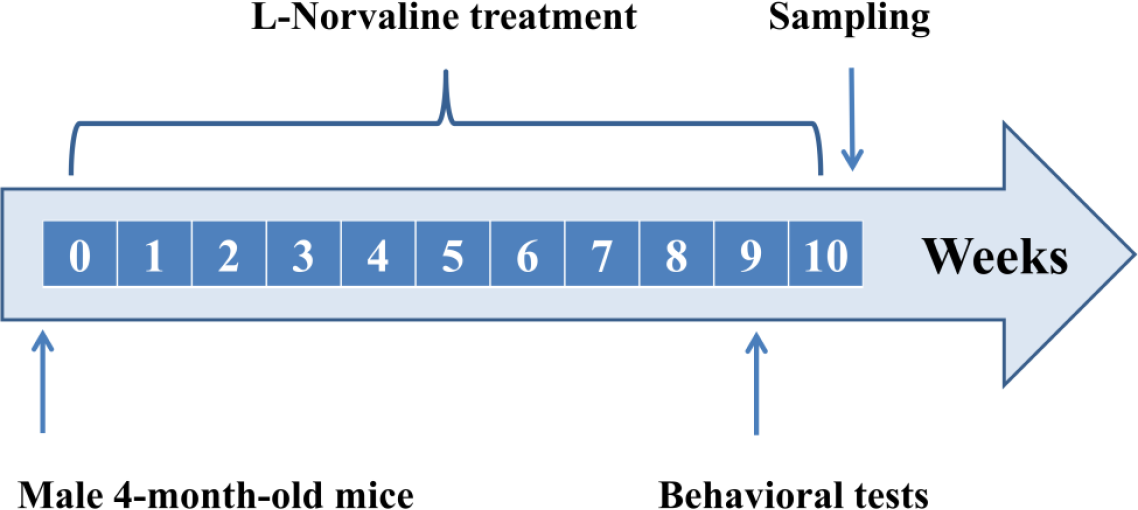
Experimental design.

**Figure S2.**
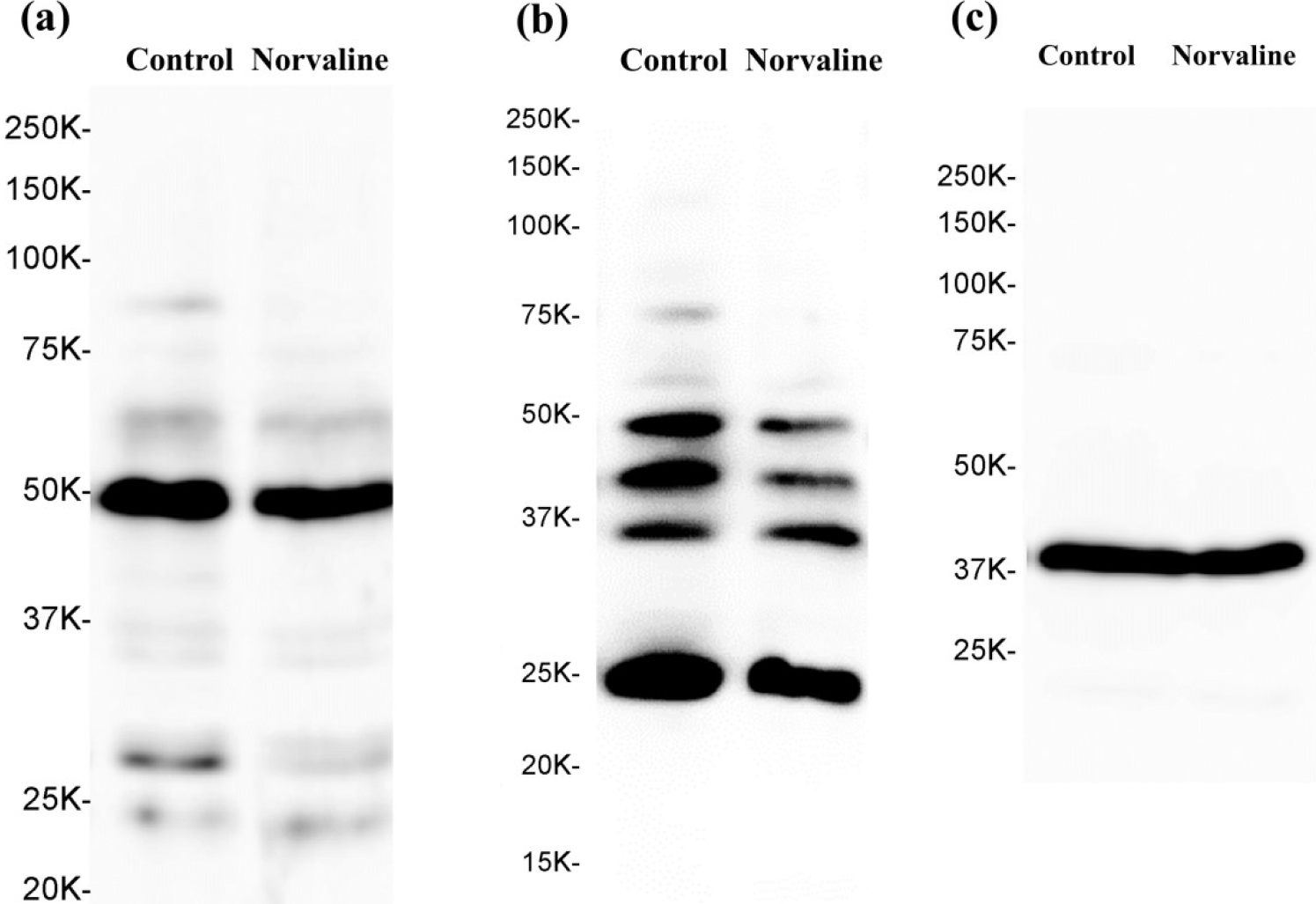
Western blots of A11 (a), OC (b), and β-actin.

**Table S1.** KCPS Primary Antibody Selection.

| Ab Code | Type | Protein Name | Host Species | MW Range | Dilution | Ab Conc'n |
| --- | --- | --- | --- | --- | --- | --- |
| NK178 | pan-specific | TrkA | RpAb | 87 kDa | 1 : 250 | 4 µg/ml |
| NN031 | pan-specific | Cyclin E | MmAb | 47 kDa | 1 : 500 | 2 µg/ml |
| NP033 | pan-specific | PP2A B' (B56) | RpAb | 56 kDa | 1 : 3000 | unknown conc'n |
| NN171-2 | pan-specific | Synapsin 1 isoform 1a | MmAb | 74 kDa | 1 : 1000 | 1 µg/ml |
| NN294-1 | pan-specific | Neurologin-1 | MmAb | 120 kDa | 1 : 250 | 4 µg/ml |
| NN362-1 | pan-specific | Vesicular glutamate transporter 3 | MmAb | 65 kDa | 1 : 250 | 4 µg/ml |
| N370-1 | pan-specific | Synaptophysin | MmAb | 38 kDa | 1 : 250 | 4 µg/ml |
| NN361-1 | pan-specific | Vesicular glutamate transporter 1 | MmAb | 60 kDa | 1 : 250 | 4 µg/ml |
| NN350-1 | pan-specific | Short transient receptor potential | MmAb | 110 kDa | 1 : 250 | 4 µg/ml |
| NN344-1 | pan-specific | Synaptotagmin-10 | MmAb | 60 kDa | 1 : 250 | 4 µg/ml |
| NN244-2 | pan-specific | Proto-oncogene tyrosine-protein | RpAb | 140 kDa | 1 : 250 | 4 µg/ml |
| NN367-1 | pan-specific | Synaptotagmin-6 | MmAb | 55 kDa | 1 : 250 | 4 µg/ml |
| NN228-1 | pan-specific | Voltage-dependent L-type calcium | MmAb | 78 kDa | 1 : 250 | 4 µg/ml |
| NN345-1 | pan-specific | Synaptotagmin-12 | MmAb | 45 kDa | 1 : 250 | 4 µg/ml |
| CN0010-1 | pan-specific | b-Actin | RpAb | 42 kDa | 1 : 1000 | 1 µg/ml |
| NN194-1 | pan-specific | Amyloid Fibrils (OC) | RpAb | 50 +Fragments | 1:1000 | 1 µg/ml |
| NN198-1 | pan-specific | Amyloid Oligomers (A11) | RpAb | 50 +Fragments | 1:1000 | 1 µg/ml |
| CN001-1 | pan-specific | Beta-Actin | RpAb | 42 kDa | 1:1000 | 1 µg/ml |

**Table S2.**
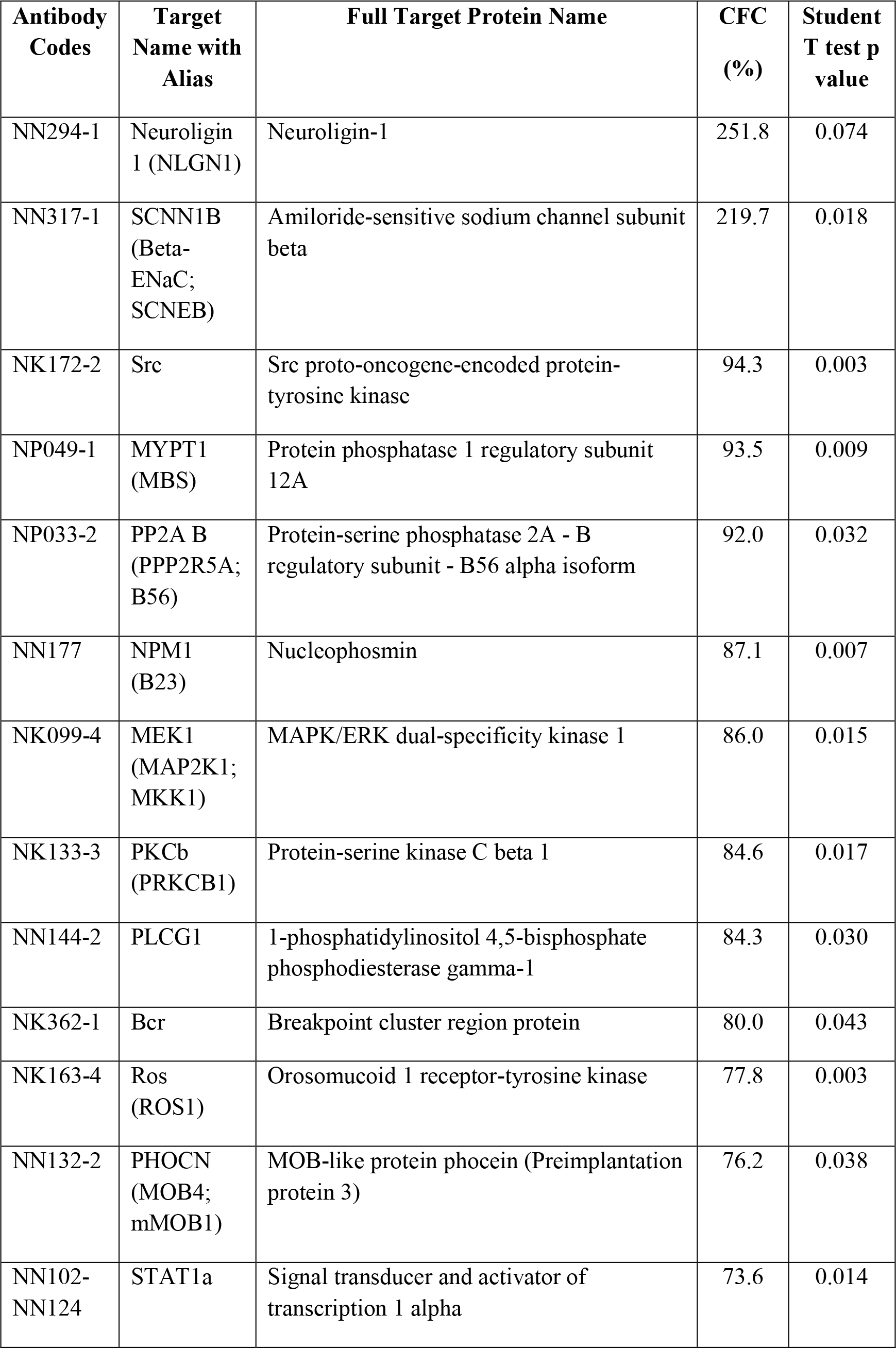
Selected results of the antibody array. Only proteins with significant (p<0.05) fold change from control (CFC) are displayed (except for Neuroligin with 252% of change). The cut-off is set at ±19% change.

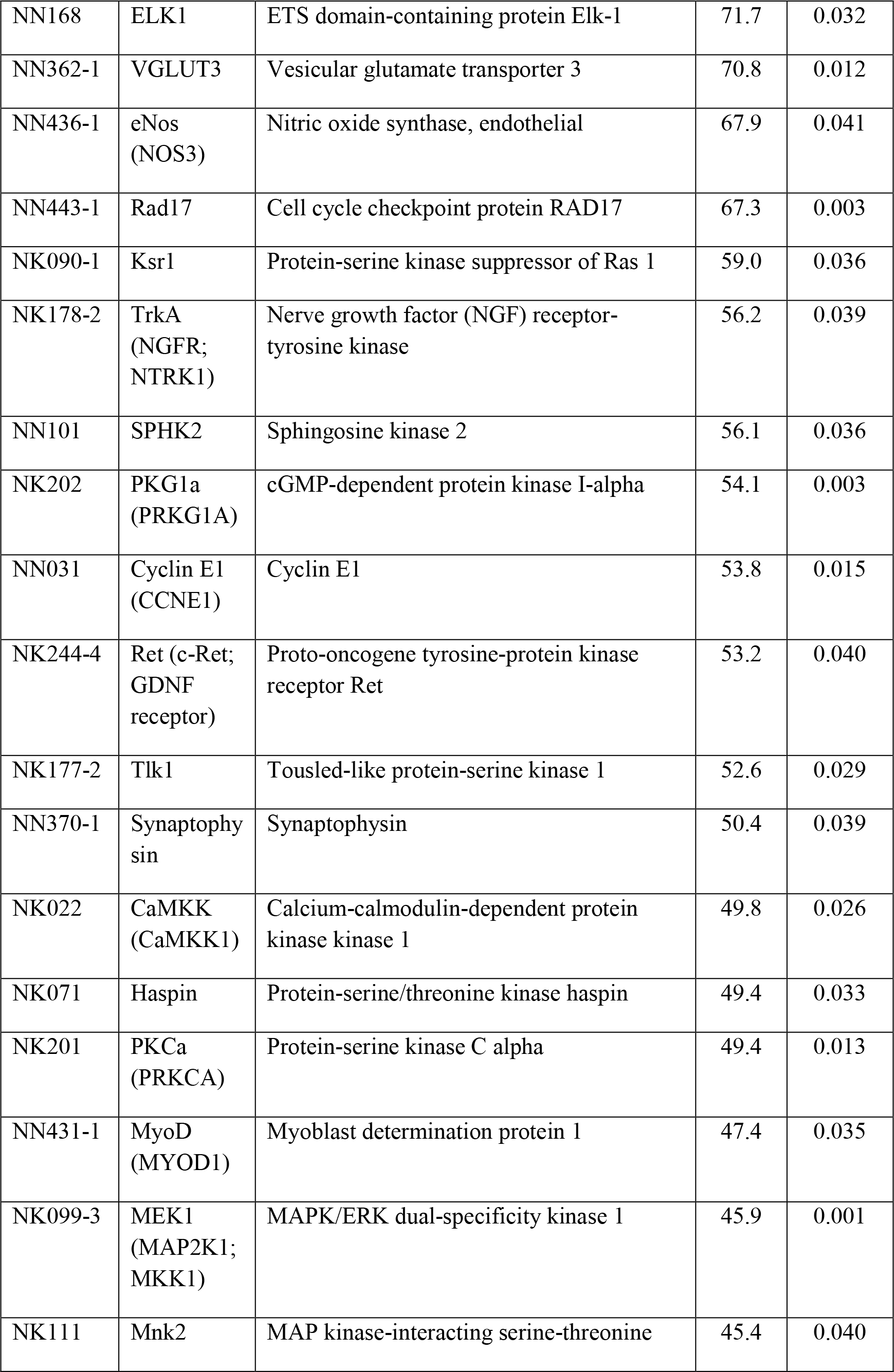

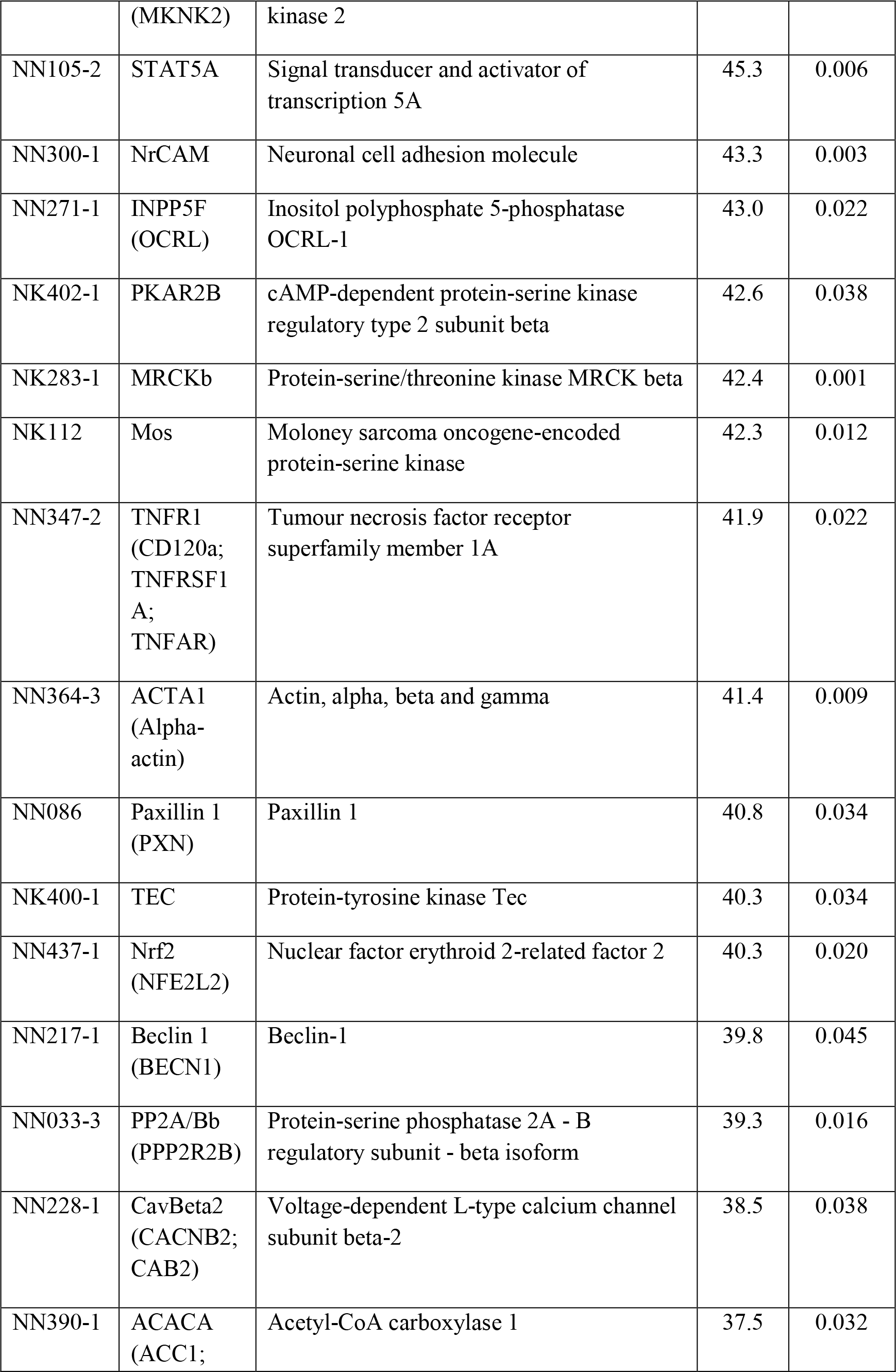

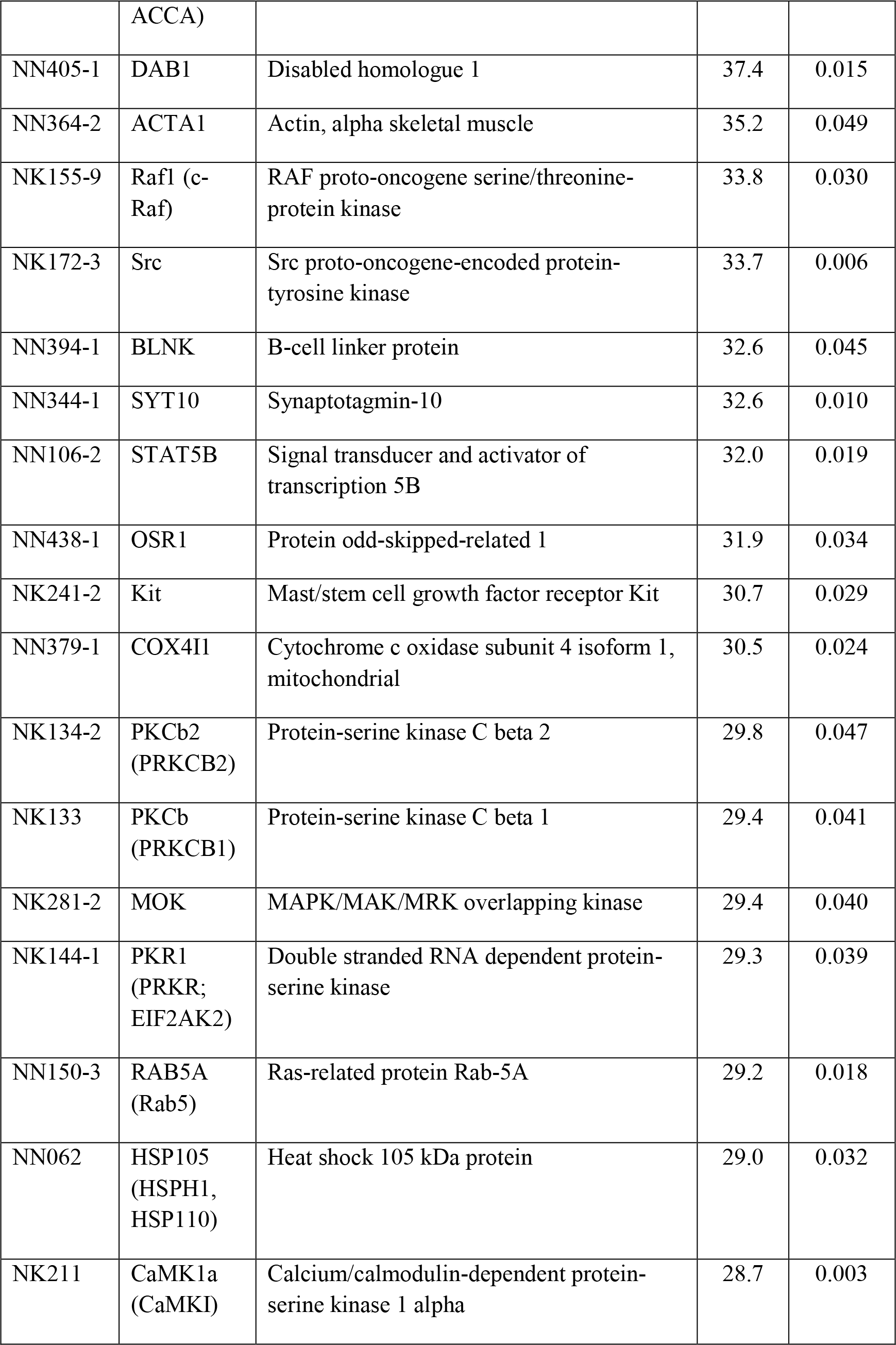

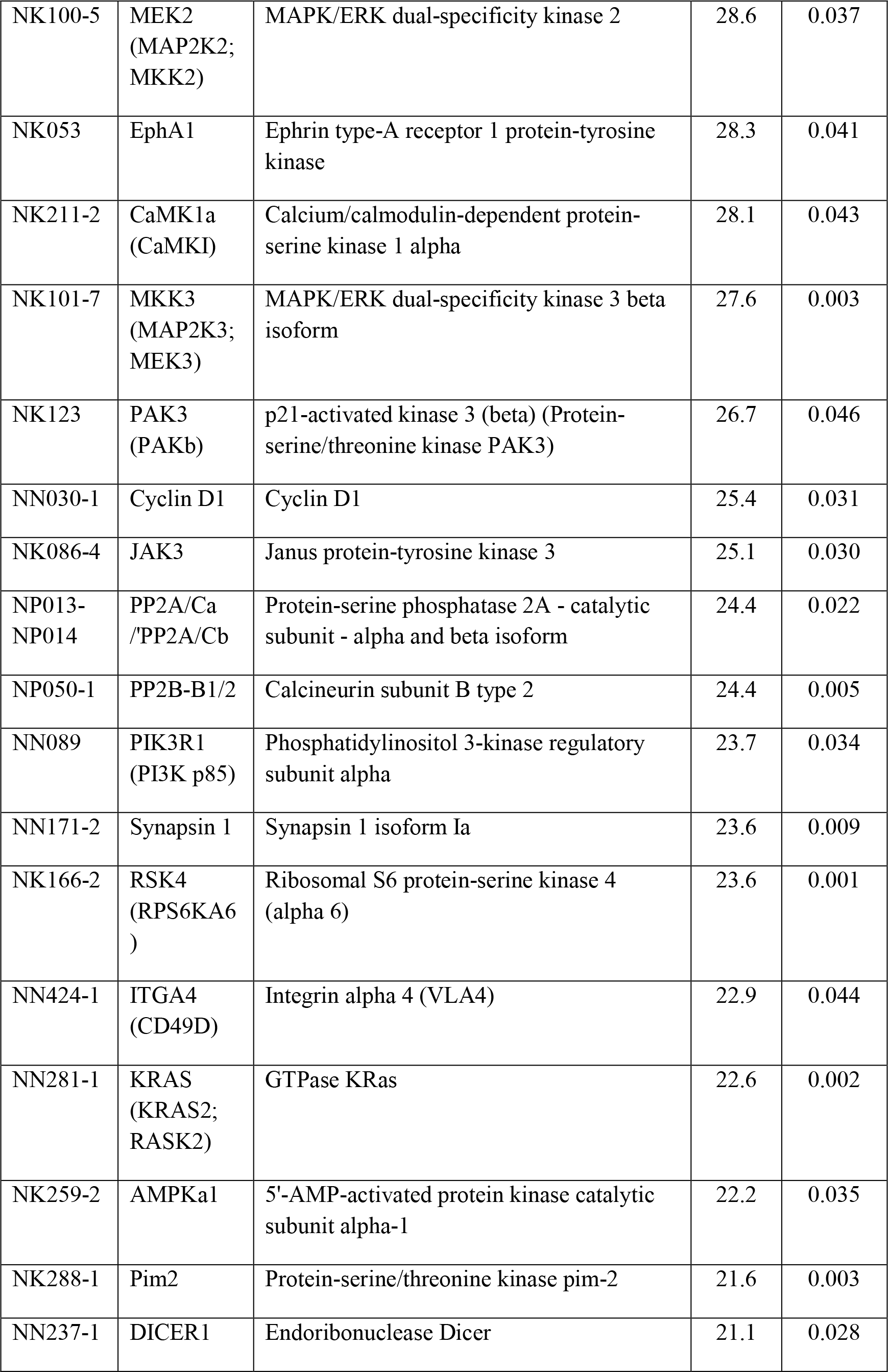

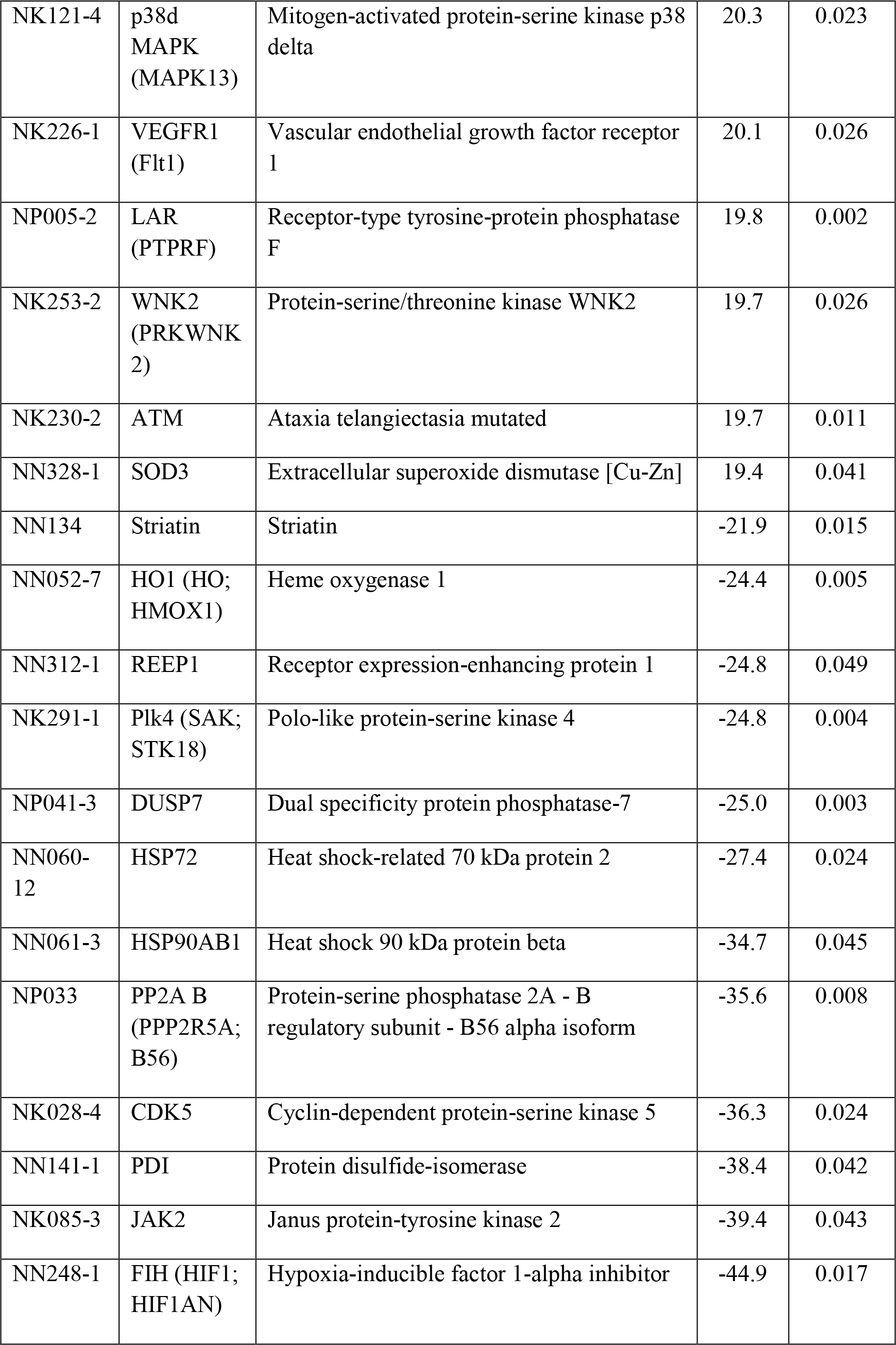

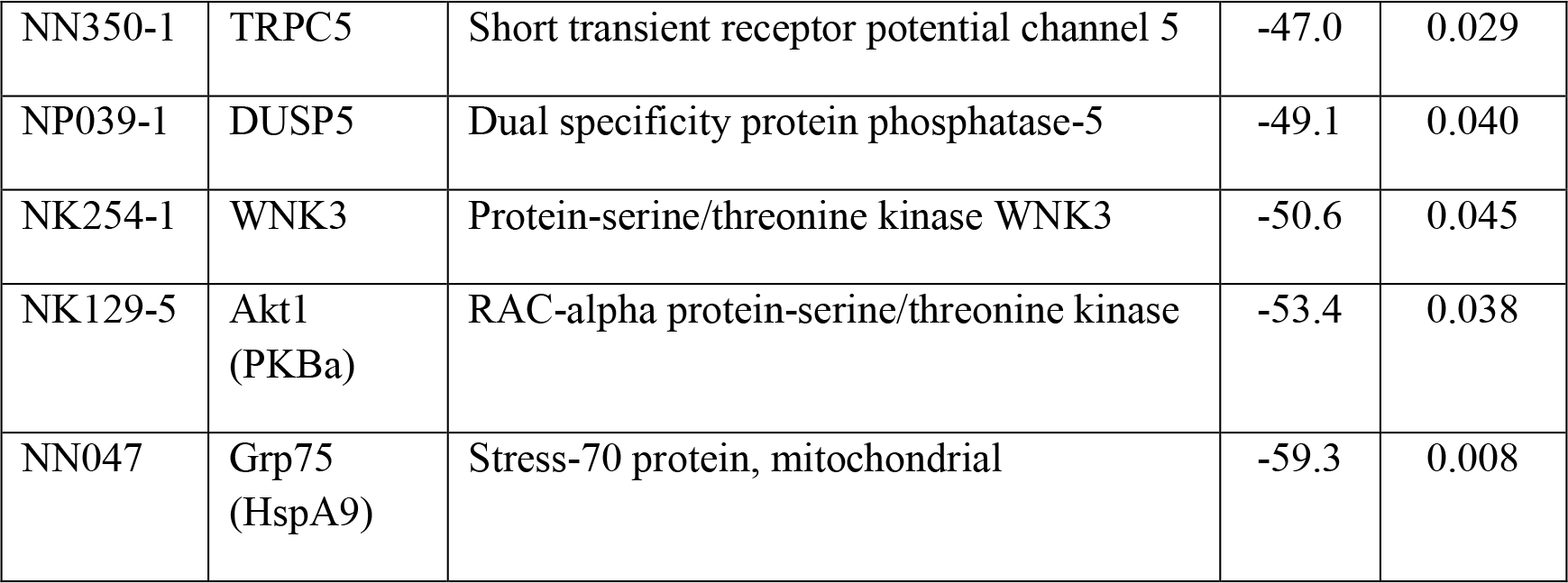

**Figure S3.**
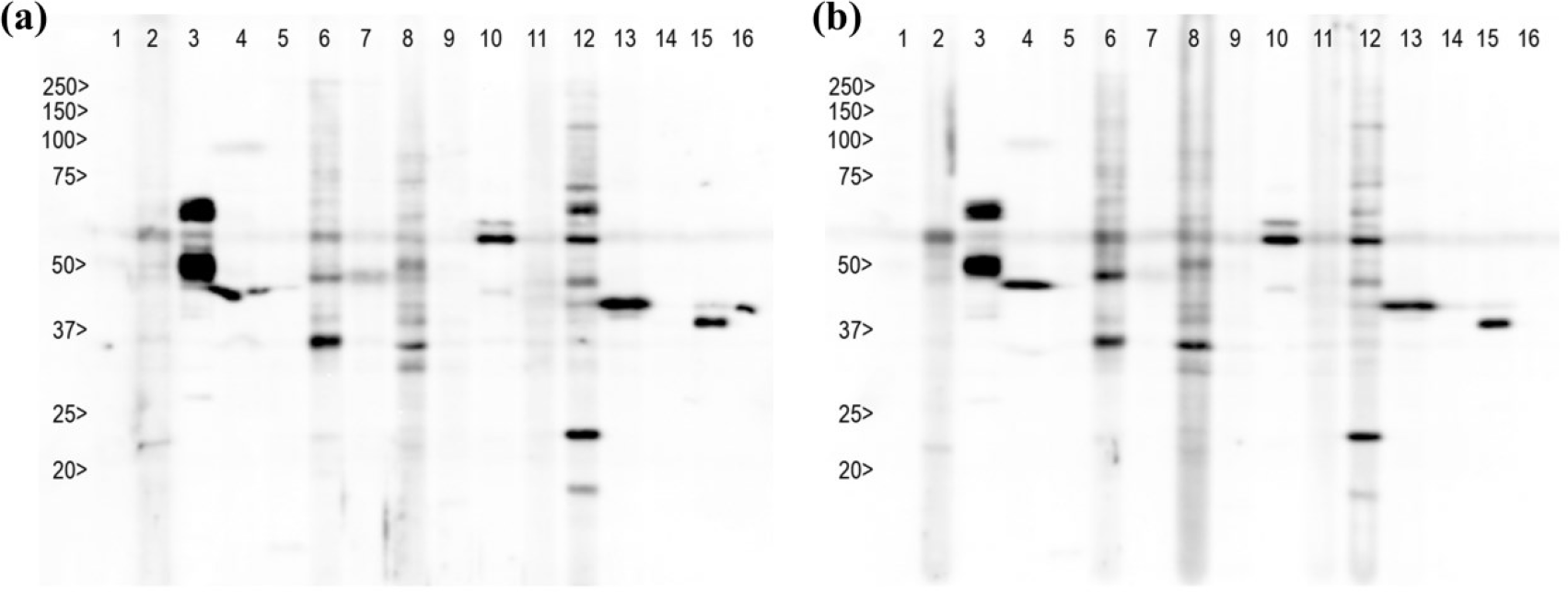
KCPS Western blots.(a) Control. (b) Treatment. Lane 2-TrkA, lane 3-Cyclin E, lane 4-PP2A B′ (B56), lane 5-Synapsin 1 isoform 1a,lane 6-Neuroligin-1, lane 7-Vesicular glutamate transporter 3, lane 8-Synaptophysin, lane 9-Vesicular glutamate transporter 1, lane 10-Short transient receptor potential, lane 11-Synaptotagmin-10, lane 12-Proto-oncogene tyrosine-protein, lane 13-Synaptotagmin-6, lane 14-Voltage-dependent L-type calcium channel, lane 15-Synaptotagmin-12, lane16-β-Actin.

## Author contributions

Baruh Polis and Abraham Samson designed experiments. Baruh Polis and Kolluru Devi Dutt Srikanth performed experiments and analyzed the data. Hava Gil-Henn advised and supervised experiments. Evan Elliott supervised parts of the experiments. Baruh Polis wrote the manuscript and Abraham Samson, Hava Gil-Henn, Evan Elliott edited the manuscript.

## Acknowledgments

This research was supported by a Marie Curie CIG grant 322113, a Leir foundation grant, a Ginzburg family foundation grant, and a Katz foundation grant to AOS. We gratefully acknowledge Mr. Basem Hijazi for his valuable advice in the statistical analysis and Dr. Zohar Gavish for his help with immunohistochemistry.

